# A bottom-up approach identifies the antipsychotic and antineoplastic trifluoperazine and the ribose derivative deoxytubercidin as novel microglial phagocytosis inhibitors

**DOI:** 10.1101/2024.06.17.599284

**Authors:** Noelia Rodriguez-Iglesias, Iñaki Paris, Jorge Valero, Lorena Cañas-Zabala, Alejandro Carretero, Klas Hatje, Jitao David Zhang, Christoph Patsch, Markus Britschgi, Simon Gutbier, Amanda Sierra

## Abstract

Phagocytosis is an indispensable function of microglia, the brain professional phagocytes. Microglia are particularly efficient phagocytosing cells that undergo programmed cell death (apoptosis) in physiological conditions. However, mounting evidence suggests microglial phagocytosis dysfunction in multiple brain disorders. These observations prompted us to search for phagocytosis modulators (enhancers or inhibitors) with therapeutic potential. We used a bottom-up strategy that consisted on the identification of phagocytosis modulators using phenotypic high throughput screenings (HTSs) in cell culture and validation in organotypic cultures and *in vivo*. We performed two complementary HTS campagnes: at Achucarro, we used primary cultures of mouse microglia and compounds of the Prestwick Chemical Library; at Roche, we used human iPSC derived macrophage-like cells and a proprietary chemo-genomic library with 2,200 compounds with known mechanism-of-action. Next, we validated the more robust compounds using hippocampal organotypic cultures and identified two hits: trifluoperazine, a dopaminergic and adrenergic antagonist used as an antipsychotic and antineoplastic; and deoxytubercidin, a ribose derivative. Finally, we tested whether these compounds were able to modulate phagocytosis of apoptotic newborn cells in the adult hippocampal neurogenic niche *in vivo* by administering them into the mouse hippocampus using osmotic minipumps. We confirmed that both trifluoperazine and deoxytubercidin have anti-phagocytic activity *in vivo*, and validated our bottom-up strategy to identify novel phagocytosis modulators. These results show that chemical libraries with anotated mechanism of action are an starting point for the pharmacological modulation of microglia in drug discovery projects aiming at the therapeutic manipulation of phagocytosis in brain diseases.

**GRAPHICAL ABSTRACT:** 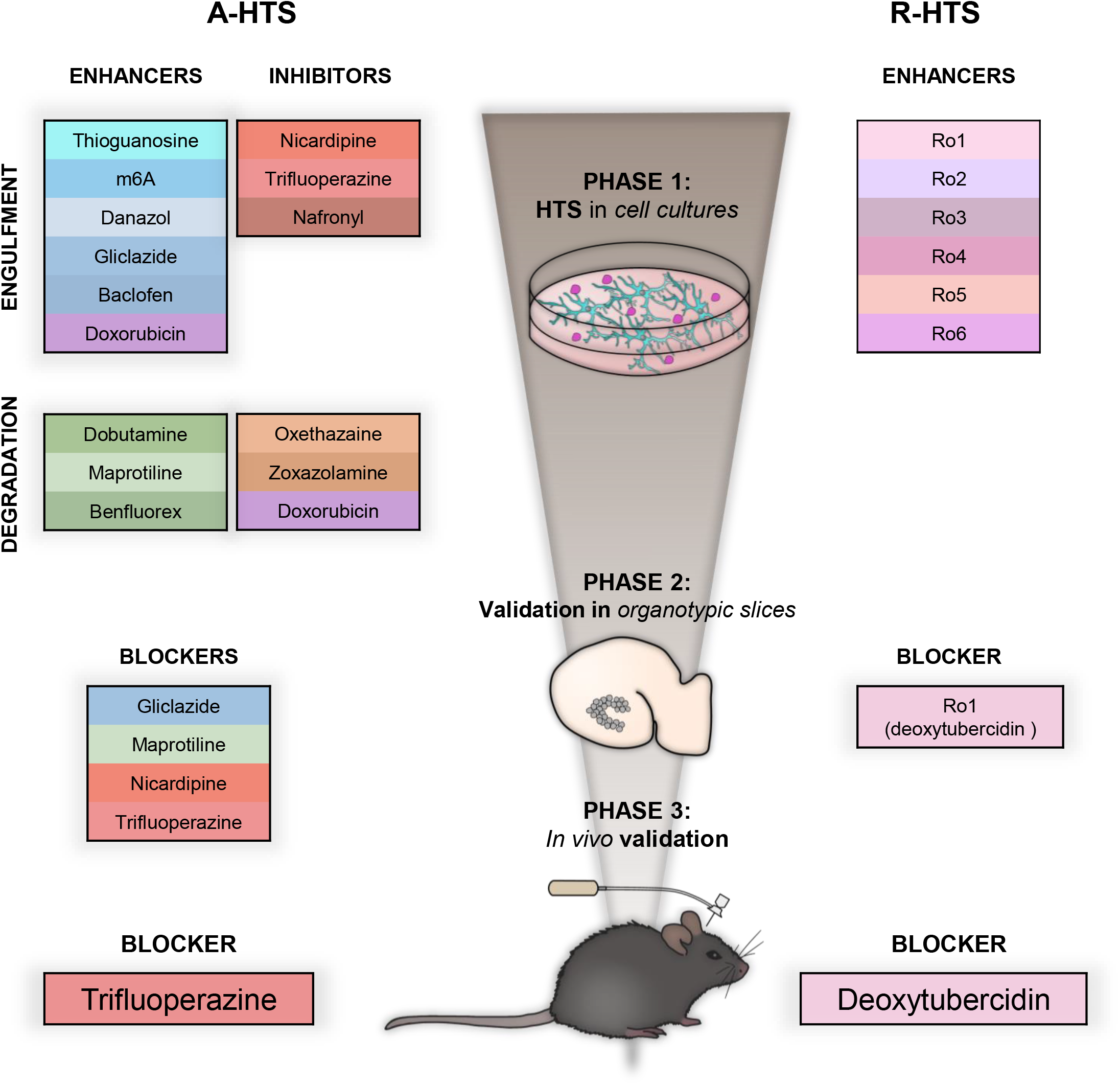

## INTRODUCTION

Phagocytosis, described by Metchnikoff late in the 19th century, refers to the process of detection, engulfment, and degradation of large particles (>0.5μm), which include dead cells, microbes, and extracellular debris (Mehrotra and Ravichandran, 2022). In the brain, phagocytosis is largely executed by the resident innate immune cells of the brain parenchyma, the microglia (Sierra et al., 2013). Equipped with motile processes (Nimmerjahn et al., 2005) and an array of surface receptors comprising the “sensome” (Hickman et al., 2013), microglia are very efficient phagocytes which can target cells that undergo programmed cell death or apoptosis, a type of phagocytosis named efferocytosis. During this process, engulfment is tightly controlled by overlapping “find-me”, “eat-me”, and “dońt-eat-me” signals that ensure the fast and specific recognition of the apoptotic cargo (Mehrotra and Ravichandran, 2022). For example, under physiological conditions, they engulf the excess of newborn cells produced in neurogenic niches (Sierra et al., 2010; Diaz-Aparicio et al., 2020) and degrade them in few hours (Kamei and Okabe, 2023). However, in some pathological conditions, the signaling responsible for the coupling between apoptosis and phagocytois is disrupted and phagocytosis efficiency is lost.

Phagocytosis can be impaired in conditions such as epilepsy (Abiega et al., 2016; Sierra□Torre et al., 2020) and stroke (Beccari et al., 2023) through multiple mechanisms that include reduced access to oxygen and nutrients, resulting in stalled microglial processes (Beccari et al., 2023); or widespread release of the neurotransmitter and “find-me” signal ATP, turning microglia “blinded” to the presence of apoptotic cells (Abiega et al., 2016). Failure to engulf apoptotic cells leads to delayed clearance and their accumulation in the parenchyma. Non-phagocytosed apoptotic cells will release toxic intracellular components that could act as danger-associated molecular patterns (DAMPS) and mount inflammatory responses (Kourtzelis et al., 2020), adding a secondary damage to the initial insult. In addition to impaired recognition of apoptotic cells, phagocytosis may also become dysfunctional as a result of misrecognizing viable cells, leading to their demise, a process termed phagoptosis (Brown and Neher, 2014). Under some inflammatory and pathological conditions, such as ischemia or the presence of certain pathogenic amyloid-beta peptide species, neurons transiently expose the “eat-me” signal phosphatydilserine (PS), triggering engulfing and execution of cell death by microglia (Brown, Guy C, 2024). Together, these observations in clinic and preclinical models suggest that targeting microglial phagocytosis with enhancers or activators in diseases where it is defective, or inhibitors in diseases where it is exacerbated could have therapeutic value to accelerate the clearance and recovery of the brain parenchyma in a large number of brain disorders. One well-known phagocytosis inhibitor is clopidogrel, an antiplatelet agent used to prevent blood clots that antagonises P2RY12 (Sipe et al., 2016), a purinergic receptor which is used by microglia to recognize apoptotic cells (Diaz-Aparicio et al., 2020). In addition, there are ongoing clinical trials targeting sialic acid-binding immunoglobulin-like lectins (SIGLECs) (Wißfeld et al., 2024) or triggering receptor expressed in myeloid cells 2 (TREM2)(Paul et al., 2021), both of which are involved in phagocytosis. However, there are no current approved therapies aimed at modulating microglial phagocytosis.

To address this gap and identify novel pathways modulating microglial phagocytosis efficiency, here we have used a bottom-up approach to identify compounds that could modulate microglial phagocytosis efficiency, from cell cultures to *in vivo*. We used two complementary high throughput phenotypic screenings: at Achucarro (A-HTS), primary microglia were screened 600 compound of the Prestwick Chemical Library; at Roche (R-HTS) we employed human iPSC-derived macrophage-like cells which are compromised in phagocytosis due to increased expression of alpha-synuclein that is caused by an allelic triplication of the *SNCA* gene (3xSNCA) (Haenseler et al., 2017) and screened with Rochés proprietary chemogenomic library (Roudnicky et al., 2020). We initially identified 12 A-HTS compounds and 6 R-HTS compounds with phagocytosis modulatory potential that were then validated in organotypic hippocampal cultures and in mice. This approach allowed us to identify the dopaminergic/α- adrenergic antagonist trifluoperazine and the ribose derivative deoxytubercidin as inhibitors of microglial phagocytosis. The pathways in which these compounds are active can now be explored for their therapeutic potential to treat diseases related to indiscriminate phagocytosis.

## METHODS

### Mice

2-month-old fms-EGFP (MacGreen) mice in which microglia express the green fluorescent reporter under the expression of the fms promoter (Sasmono et al., 2003; Sierra et al., 2007) were used. Mice were housed in 12:12 h light cycle with ad libitum access to food and water. Both males and females were used and pooled together, unless otherwise noted. All procedures followed the European Directive 2010/63/EU and were approved by the Ethics Committees of the University of the Basque Country EHU/UPV (Leioa, Spain; M20-2019-068).

### Mouse primary microglia culture

Primary microglia cultures were performed as previously described (Beccari et al., 2018). Briefly, P0-P1 fms-EGFP mice brains were carefully placed in Hank’s balanced salt solution (HBSS, Hyclone) filled Petri dishes under a magnifying scope, meninges were removed, and the brains enzymatically digested with papain (20 U/ml, Sigma-Aldrich) and DNAse (150 U/μl, Invitrogen) for 15 min at 37°C. The homogenization process was helped by mechanical disaggregation by carefully pipetting. The resulting cell suspension was filtered through a 40μm cell strainer (Fisher) into a 50 ml Falcon tube containing 10 ml of 20% heat-inactivated Fetal Bovine Serum (FBS; Invitrogen) in HBSS to stop the enzymatic reactions. Afterward, the cell suspension was pelleted at 200xG for 5 min, and the pellet was resuspended in 1 ml Dulbecco’s Modified Eagle’s Medium (DMEM, Invitrogen) supplemented with 10% fetal bovine serum (FBS) and 1% Penicillin-Streptomycin (Invitrogen). Cells from two brains were seeded in a Poly-L-Lysine-coated (15 μl/ml, Sigma-Aldrich) culture flask. The medium was changed every 3–4 d and enriched with granulocyte-macrophage colony-stimulating factor (5 ng/ml GM-CSF, Sigma-Aldrich). After approximately 14 d at 37°C and 5% CO_2_, microglia were harvested by shaking at 100 rpm, 37°C, 4 h.

### SH-SY5Y vampire culture

SH-SY5Y-Vampire (tFP602, a red-shifted version of turbo RFP; REF: P20303, Innoprot) cells were routinely maintained in DMEM supplemented with 10% FBS and 1% Penicillin-Streptomycin. Media was changed every 2 d until reaching confluence. Once confluent cells were passaged using Trypsin/EDTA for 5 min at 37°C and 5% CO_2_, the cell suspension was then mixed with DMEM to stop the enzymatic reaction and transferred to a 50 ml falcon tube. The cells were then pelleted at 200xG and reseeded.

### A-HTS Screening

Microglia were seeded at 30,000 cells/well in 96 well black-wall, clear-bottom imaging plates (BD) 24 h prior to the phagocytosis assay. For the phagocytosis assay, we used the method previously described by our group (Beccari et al., 2018). Briefly, SH-SY5Y vampire cells were treated with 3 µM staurosporine for 3 h in an incubator at 37°C and 5% CO_2_. Dead cells were harvested by collecting the supernatant in 50 ml falcon tubes and pelleted at 200xG for 5 min, the supernatants were discarded, and the pelleted cells were carefully resuspended in media. Apoptotic cells were counted using a trypan blue solution and a Neubauer cell counting chamber and were then fed to microglia in a 1:1 ratio (30,000 apoptotic cells/well).

A chemical library comprising 600 compounds was sourced from the Prestwick Chemical Library, based on the compounds’ chemical and pharmacological diversity, bioavailability, and safety for humans. Compounds were diluted to working concentration using the Hamilton’s Microlab Star automated liquid handling workstation, and the diluted compounds were manually dispensed to the cells. The compound screening was performed in 2 phases: in the first phase, we tested the 600 compounds in three replicates at 10 μM and selected the top 30 hit compounds based on their effects on engulfment and degradation of apoptotic cells. To determine which compounds were of interest in the first phase, we set a threshold equal to the mean activity of all compounds ± standard deviation. In the second phase, we re-screened the 30 hit compounds at 3 different concentrations (1, 10, and 100 μm, three replicas per concentration).

In each of the phases, we discriminated the effects of the compounds on either engulfment or degradation of apoptotic cells by using two administration paradigms. To check the effects on engulfment, microglia were incubated with the compounds for 2 hours prior to co-incubation with the apoptotic cells for 1 additional hour while maintaining the compounds in the medium. To check the effects on degradation, microglia were co-incubated with the apoptotic cells for 1 hour and, after removing the excess of un-phagocytosed apoptotic cells, they were allowed to degrade the engulfed cells in the presence of fresh media with the compounds for 2 hours. After both administration paradigms, media was removed, cells were fixed with 4% formaldehyde (Sigma-Aldrich) and processed for immunofluorescence in the wells. Shortly, they were incubated with blocking solution (Triton-X100 (Sigma-Aldrich) 0,1% and bovine serum albumin (BSA, Sigma-Aldrich) 0,5% in phosphate-buffered saline (PBS, Sigma-Aldrich) for 1 hour at room temperature. The blocking period was followed by the primary antibody solution with chicken-anti-GFP antibody (Aves Laboratories, Tigard, OR) at 1:1000 dilution in blocking solution for 1 hour. After the incubation, the cells were subsequently washed with PBS, and incubated for 1 hour in the secondary antibody solution, which contains blocking solution + Goat-anti-Chicken-Alexa-488 conjugated antibody (Life Technologies) at a 1:500 dilution + 4’,6-diamidino-2-phenylindole (DAPI, Sigma-Aldrich) at a 1:1000 dilution.

### A-HTS hit selection and confirmation

The following variables for each compound were obtained: number of microglia, mean area of Vampire^+^ particles per microglia, percentage of phagocytic microglia, and number of phagocytosed objects per microglia. To decide which variable was better suited to study microglial phagocytosis activity in this dataset, we performed principal component analysis (PCA) for both engulfment and degradation. We visualized the effect of the compounds using a scatter plot to compare PC1 and PC2 and observed that most compounds clustered homogenously. However, some compounds did not cluster with the rest of the compounds, indicating that they were some potential candidate compounds to modulate phagocytosis (effects if engulfment displayed in **Supp. Fig. 1A**). We then analyzed how each principal component (PC) separated each compound and the way each variable weighted in each PC: PC1 explained close to 60% of the variance, as it had the largest eigenvalue, whereas PC2 explained 32% (**Supp. Fig. 1B**). The variables that contributed the most (had the highest % of loadings) to both PC1 and PC2 were the number of phagocytosed objects per microglia and the number of microglia (**Supp. Fig. 1C, D**). Thus, we selected the variable number of phagocytosed particles per microglia to assess the potential of these compounds to enhance or block either engulfment or degradation.

To reduce the variability between technical replicates (wells), we filtered out those that displayed large variability (standard deviation, SD) in the number of microglia (SD>30% of mean) and in the number of phagocytosed particles per microglia (SD>30% of mean). After this 2-step filtering procedure, we checked the integrity of the data by analyzing two parameters in the dataset: 1, the variability between compounds, to determine how similar was the effect of all compounds by comparing the effect of each compound *vs* the mean effect of all compounds; and 2, the variability between replicas, to determine how similar is the variability of the effect of all compounds by comparing the SD (between wells) of each compound *vs* the mean SD of all compounds. The filtering procedure had no effect on the variability between compounds but reduced the variability between replicas (P<0.001, **Supp. Fig. 1E-G**). Thus, variability was reduced while preserving the differences between compounds for both engulfment and degradation. Next, we removed the data of those compounds that had a toxic effect on microglia (<70% of the control). From 600 initial compounds, 63 were eliminated due to their toxicity in the engulfment experiments, 23 were eliminated due to their toxicity during the degradation experiments, and 4 were eliminated due to their toxicity in both series of experiments (leaving a final number of 510 compounds). This final filtering did not affect either the variability between replicas or the variability between compounds (**Supp. Fig. 1H,I**).

The same analysis was performed during Phase 2. PCA showed that while most compounds clustered homogenously, some compounds did not cluster along PC 1, indicating potential phagocytosis modulators (**Supp. Fig. 2A**). PC1 explained about 71% of the variance, and the variable that contributed the most was the number of phagocytosed objects per microglia (**Supp**. **Fig. 2B, C**). PC2 represented about 23% of the variance, and the variable that contributed the most was the number of microglia (**Supp**. **Fig. 2B, D**). We followed the same data homogenization strategy as in Phase 1 to decrease variability between replicas by filtering out replicas where the number of microglia and the number of phagocytosed objects showed high variability between wells, and finally filtering out compounds based on their toxicity (**Supp. Fig. 2E-H**). As in the first phase, we achieved a decrease in the variability between replicas (P<0.001 in engulfment, no significant differences in degradation) without affecting variability between compounds (P>0.05) (**Supp. Fig. 2E, F**). In this phase, no compounds were eliminated due to toxicity on microglia: only three compounds at their highest dose (100μM) showed toxic effects on microglia and were subsequently removed (at that dose only). The variability between replicas and compounds was also monitored after eliminating toxic compounds. The z-score that summarized the effect of each compound across the three concentrations was calculated as follows:

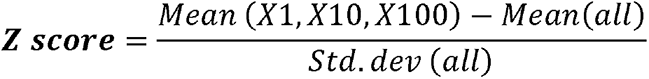

### Human iPSC culture and differentiation to macrophage-like cells

Two human iPSC lines were used: SBNeo1 as control and SFC831-03-03 carrying a synuclein triplication (3xSNCA), both generated in the StemBANCC consortia and deposited at European Bank for iPSCs) (Gutbier et al., 2020). A single cell suspension of iPSCs (4×10^6^ cells) was plated into AggreWell 800 iPSCs were plated into AggreWell 800 (StemCell Technologies, Vancouver, BC, Canada) plates in mTesR1 supplemented with 10 μM ROCK inhibitor (Y27632, Calbiochem/ Millipore, Burlington, MA, USA). Next, cells were treated with 50 ng/mL human bone morphogenetic protein 4 (hBMP4) (R&D Systems, Minneapolis, Minnesota, USA), 50 ng/mL human vascular endothelial growth factor (hVEGF) (R&D Systems), and 20 ng/mL human stem cell factor (hSCF) (R&D Systems). After 4 days, embryonic bodies were collected and used to generate myeloid factories by growing them in X-VIVO 15 medium (Lonza, Basel, Switzerland) supplemented with 2 mM Glutamax, 1% penicillin/streptomycin, 50 ug/mL mercaptoethanol, macrophage colony stimulating factor (M-CSF, 100 ng/mL)(Miltenyi Biotech, Bergisch Gladbach, Germany), and IL3 (25 ng/mL) (Miltenyi Biotech). Macrophage precursors were harvested from myeloid factories over aperiod of 6 weeks from a 4 layer cell disk and accumulated in suspension culture. To differentiate macrophages 8,000/ cells per well were plated in 384 well plates (Corning Cat. No. 3575, Corning Inc., New York, USA) in 20 ul XVIVO 15 media containing 100 ng/ml MCSF, after 3 days 16 ul XVIVO 15 media containing 100 ng/ml MCSF was added.

In phagocytosis experiments, neural stem cells (NSCs) derived from the iPSCs were used (Cusulin et al., 2019). Apoptosis was induced by incubation of NSC with 10 μM staurosporine for 4 hours. NSC were detached using accutase, collected by centrifugation, resuspended and washed twice in Live Cell Imaging Solution (Invitrogen, Waltham, MA, USA). Incubated for 15 min with pHrodo Red succinimidyl ester (Invitrogen), washed twice with imaging solution, fixated in 4% PFA solution, washed twice with Live cell imaging solution, pelleted and stored at -80 °C.

### R-HTS screening

At day 5 of differentiation to macrophage-like, Rochés compounds prediluted in culture media were added at 1 μM final concentration, with 6 replicate plates and incubated for 24 h. In the last two hours, thawed apoptotic phRodo-labeled NSCs (24,000/well) were added to start the phagocytosis experiment. After incubation cells were washed twice with DPBS (Gibco) and fixated for 10 min with 4% PFA containing Hoechst-33342 (Invitrogen) in DPBS. All aspiration steps were performed leaving 10 ul dead volume in the assay plate to avoid disturbance of the cell layer, dispense of solvents was performed at a dispensing ofset of 1.5 mm from assay plate bottom at a speed of 10 ul/sec, followed by a tip touch at the edge of the well to remove remaining droplets from the pipett tips. All pipetting steps were performed on the Biomek FX automated workstation (BeckmanCoulter, Pasadena, CA, USA). After fixation, cells were washed again with DPBS and imaged using operetta cls (PerkinElmer; Waltham, MA, USA) hcs system

### R-H TS hit selection and confirmation

In total 2,200 compounds were screened at 1 uM with 6 replicates for their potential to increase phagocytosis in the macrophages derived from the 3xSNCA iPSC line. The SBNeo line was used as a control. Compounds that increased phagocytosis (ratio ofphagocytic microglia relative to control>1.3) and had a low standard error of the mean (SEM<0.2) between the six replicates were picked for hit confirmation in dose response (starting at 10 μM in 1:2 dilution) in macrophages derived from iPSC of apparently healthy donor and in the iPSC line carrying the alpha synuclein triplication. Compounds that confirmed their effect in dose response were picked for testing in organotypic slices.

### Organotypic cultures and compound incubation

Hippocampal organotypic slices were prepared following the protocol described in (Beccari et al., 2018). Briefly, postnatal day 7 (PND 7) fms-eGFP mice were decapitated and their brains were extracted and placed in cold HBSS (Cytiva). Right and left hippocampi were dissected under a binocular magnifier (Olympus SZ51) and sectioned into 350 μm slices using a tissue chopper (McIlwain). Individualized slices were then transferred to 0.4 μm polytetrafluoroethylene (PTFE) culture inserts (Merck), which were placed in six-well plates. Each well contained 1ml of organotypic culture medium that consisted of 50% Neurobasal medium (Gibco) supplemented with 0.5% B27 (Gibco), 25% horse serum (Ibian Technologies), 1% Glutamax (Gibco), 1% penicillin/streptomycin (Fisher), 1% glucose solution in MilliQ water (Gibco), and HBSS. Hippocampal slices were kept in culture at 37°C and 5% CO_2_ for 7 days until the day of the experiment, and the medium was changed the day after the culture and every 2 days. At day 7of culture, hippocampal slices were incubated with 100 μM of the compounds diluted in organotypic culture medium for a total of 6 hours. After 5 hours of incubation, the medium was replaced with fresh medium containing the compounds and 5 μg/ml propidium iodide (PI). After 1 hour of incubation with compounds and PI, slices were fixed in cold 4% paraformaldehyde solution for 40 minutes and, after rinsing with 1X PBS, were stored at 4°C until immunofluorescence processing.

### Compound administration *in vivo*

Compound administration *in vivo* to 2-month-old fms-eGFP mice was carried out following the protocol described in (Diaz-Aparicio et al., 2020) with minor modifications. Briefly, osmotic minipumps (flow rate 1μl/h; Model 1003D, Alzet) and brain infusion catheter tubes (Alzet) were filled with 100 μl of dimethyl sulfoxide (DMSO; Thermo Fisher), Roche compound #1, Trifluoperazine (Selleckchem), and PSB-0739 (Tocris Bioscience) (all at 10μm concentration in PBS with 1:1000 DMSO) and connected. Pumps were incubated overnight at 37°C submerged in PBS before the surgery. Mice were anesthetized intraperitoneally with ketamine/xylazine (10/1mg/kg) and received a single dose of the analgesic buprenorphine (1mg/kg) subcutaneously. The infusion cannulae were inserted from bregma at: anterioposterior -1.7mm; laterolateral: -1.6mm; and dorsoventral -1.9mm. To avoid the removal of the cannulae, a surface of dental cement was created from the cannula to the screw. Finally, the pumps were inserted inside the back skin of the mice. After 2d, mice were transcardially perfused.

### Immunofluorescence

Immunostaining was performed following standard procedures (Beccari et al., 2018). Free-floating organotypic slices or vibratome sections were incubated in permeabilization solution (0.2% Triton-X100, 3% BSA in PBS; Sigma) for 1 hour at room temperature. Then, organotypic slices or tissue sections were incubated overnight with the primary antibodies (chicken anti-GFP, 1:1000) diluted in the permeabilization solution at 4°C. After washing with PBS, organotypic slices or tissue sections were incubated with fluorochrome-conjugated secondary antibodies (Alexa fluor 488 donkey anti-chicken; Jackson ImmunoResearch) and DAPI (5mg/ml; Sigma) diluted in the permeabilization solution for 2 hours at room temperature. After washing with PBS, tissue sections and organotypic slices were mounted on glass slides with DakoCytomation Fluorescent Mounting Medium (Agilent).

### Imaging

In the A-HTS, we used a BD pathway 855 high content automated screener. The resulting image for each well was obtained by combining 9 images that were taken as a 3 x 3 single-plane tile using an Olympus 40X dry objective and the confocal mode of the screener. The excitation and emission filters used were 380/10 and 450/10 for DAPI; 488/10 and 540/20 for GFP; and 548/20 and 570LP for Vampire. In the R-HTS, Images (9 fields per well, single z-plain, widefield, 2×2 binning) were aquired using Operetta CLS high content imaging system (PerkinElmer; Waltham, MA, USA) using a 20x air objective and Hoechst and Alexa555 predefined filter sets. Exposure was set on several plates to be within the first 3rd of the greyscale to avoid saturation. For organotypic cultures or brain sections, fluorescence immunostaining images were collected using a Leica SP8 or Leica Stellaris laser-scanning microscope using a 40x (brain tissue sections) or 63x (organotypic slices) oil-immersion objectives and a z-step of 0.7μm. All images were imported into ImageJ (FIJI) software in tiff format. For brain tissue sections, 2 to 4 15-30μm-thick z-stacks located at random positions containing DG were collected per hippocampal section, and a minimum of 5-6 coronal sections per series were analyzed. In organotypic slices, 2-5 images of the DG were acquired per slice, and each experimental condition consisted of 2-5 slices.

### Image Analysis

In A-HTS, the software BD AttoVision 1.6/855 was used for the analysis of microglial phagocytic activity in primary cultures. We segmented the images to generate regions of interest (ROI) representing the surface area of each cell. First, we segmented the nuclei based on DAPI staining and using the following parameters: shading (normalizes uneven light detection); Rb75×75 (rolling ball background subtraction); 150<pixel size (size filter: elements with an area smaller than 150 pixels were discarded). Then, we segmented each microglia cell based on GFP staining using the following parameters: close5×5 (close filter to fill in gaps in the GFP signal); Rb75×75, 100<pixel size<150,000 (elements with area smaller than 100 and larger than 150,000 pixels); dilatation width 1000 (makes objects more visible and fills in small holes in objects). Phagocytosed particles were segmented filtering the Vampire signal: Rb2×2, TopHat7×7 (an operation that extracts small elements and details from given images), 2<pixel size<50.

The parameters that we obtained were the following:

- Number of apoptotic objects per microglia cell: phagocytosed particles were counted per cell as the average Vampire^+^ particles contained within each GFP^+^ ROI.
- Phagocytosed object area: the mean area occupied by each vampire^+^ ROI that colocalized with a GFP^+^ ROI.
- Percentage of phagocytic microglia: the percentage of GFP^+^ ROIs containing at least one Vampire^+^ ROI.
- Number of microglia cells: the number of GFP+ ROIs

The data were normalized as percentages compared to the untreated control. Cellular viability was determined by counting the nuclei (DAPI). The results are expressed as the mean and standard deviation (SD). To calculate the most consistent effect (X) across concentrations (1, 10, and 100μM), we calculated a z-score from each compound:

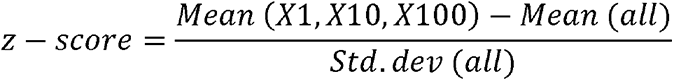

Cell death and phagocytosis analyses were carried out using unbiased stereology methods described in (Beccari et al., 2018). Apoptotic cells were determined by their nuclear morphology visualized with DAPI. These cells lost their chromatin structure and appeared condensed and/or fragmented (pyknosis/karyorrhexis). Primary and secondary necrotic cells were identified because they had incorporated propidium iodide (PI^+^ cells). Phagocytosis was defined as the formation of a 3D pouch completely surrounding an apoptotic or PI^+^ cell (Sierra et al., 2010). In organotypic slices apoptotic cells, PI-positive cells, and microglia numbers in the DG were given as a density, over a 200.000 μm3 volume (roughly, a 100 x 100 μm2 area of 20 μm of thickness). In brain tissue sections, the numbers of apoptotic and microglia cells were estimated in the volume of the DG contained in the z-stack (determined by multiplying the thickness of the stack by the area of the DG at the center of the z-stack). To obtain the absolute numbers, i.e., cells per septal hippocampus), the density values were multiplied by the volume of the septal hippocampus, which was calculated using ImageJ from images obtained with a 20x objective in an automated digital slide scanner (Pannoramic Midi II; 3D Histech Ltd, Hungary).

In R-HTS, Images were analyzed in paralell to acquisition using Harmony (PerkinElmer; Waltham, MA, USA) predifiened building blocks for detection of nuclei using the Hoechst Channel, detection of cytoplasm using the Hoechst reflection within the cytoplasm and the spot detector to detect pH-Rodo positive spots within the cytoplasm. Cells were selected as phagocytic when they had 1 or more pHrodo spots in cytoplasm. The ratio of phagocytic cells was calculated by dividing the positive cells by all valid nuclei and normalized to average of the DMSO control wells on the corresponding plate. Mean, median, SD and SEM of phagocytic index over 6 replicates was calculated and the compounds that robustly induced phagocytosis were selected for further hit validation in dose response.

### Microglia morphology analysis

Microglia morphology analysis was semi-automatically performed using an in-house built macro package for ImageJ (Cell_Shaper v5.1 https://www.achucarro.org/es/downloads/) based on the Sholl technique (Sholl, 1953). Briefly, only microglia cells located in the center of the z-stack and not touching the edges of the images were selected for the analysis in order to avoid incomplete cells. Using the first macro of the package (M1), the center and perimeter of the cell were manually selected in the z-stack images and in their maximum projection, respectively. The second macro (M2) ran automatically and used the ImageJ Sholl Analysis plugin to generate concentric spheres every 1μm starting from the selected center. The plugin calculates the number of intersections in each sphere shell for each cell, as well as the radius length of the last sphere that intersects a microglia ramification (termed here maximum length of microglia processes). The following parameters can be obtained with the macros: number of intersections per radius, sum of intersections per cell, and maximum length of the ramifications from the soma. 3D reconstructions of microglia cells were performed using the 3D viewer plugin from ImageJ.

### Statistics

For the principal component analysis PAST3 (Oslo Hammer, Ø., Harper, D.A.T., Ryan, P.D. 2001. PAST: Paleontological statistics software package for education and data analysis. Palaeontologia Electronica 4(1): 9pp. http://palaeo-electronica.org/2001_1/past/issue1_01.htm) was used. The normalized data for each variable was imported, selected, and processed using the PCA module. Between-groups comparison was performed using Sigmaplot. First, data were tested for normality and homoscedasticity. None of the data followed a normal distribution and thus were analyzed using a Kruskal–Wallis ranks test, followed by pairwise multiple comparison using Dunn’s method as a post hoc. For hippocampal cultures and *in vivo* experiments, Graphpad Prism version 10 (Graphpad Software, Boston Massachusetts, USA) was used. Normality and homoscedasticity of data were analyzed with the Kolmogorov-Smirnov test and F-test or Brown-Forsythe test, respectively. When data did not comply with these assumptions, they were transformed and analyzed using parametric tests. Two-sample experiments were analyzed using the Student’s t-test and more than two sample experiments with one-way ANOVA followed by a Dunnett’s multiple comparison test. For the correlation matrix analyses, the non-parametric Spearman’s ρ test was used to avoid the confounding effect of clustered data. Only p < 0.05 is reported to be significant except in multiple correlations, where significance was established at p < 0.01 to reduce possible random detection of significant correlations due to multiple comparisons.

## RESULTS

### Achucarro High Throughput screening (A-HTS)

In the first phase of the A-HTS, 600 Prestwick compounds were tested using 2 administration paradigms to check for effects in either engulfment or degradation, in triplicates at 10μM (**Fig. 1A, B; Supp. Fig.1**). In order to select the candidate compounds that could modulate phagocytosis, we plotted their effect on engulfment vs degradation (**Fig. 1C**), setting a threshold equal to the mean activity of all compounds ± SD. We selected the top 30 compounds that, based on the data distribution, belonged to three different groups: 1, All compounds that surpassed the threshold for enhancing engulfment and degradation; 2, those that crossed the threshold for inhibiting engulfment; and 3, an extra compound that showed a very strong enhancement of engulfment and a very strong inhibition of degradation. Those 30 candidate compounds were identified and screened in a second phase at 3 concentrations (1, 10, and 100 μM; data from one compound was lost due to technical reasons, resulting in 29 compounds tested)(**Supp. Fig. 2**). To obtain more robust data about their activity and an estimation of the working concentration of these compounds for their use *in vivo*, we analyzed them based on a z-score that accounted for a global effect of the 3 concentrations tested. Those compounds with a z-score greater than 1 or lower than -1 were selected as hits (14 compounds, of which 12 were tested later in organotypic slices)(**Fig. 1D**). Comparing the results obtained from this phase with the results obtained from phase 1, we observed that some of the compounds displayed the same effect on phase 1 and phase 2 (34.5% of compounds for engulfment and 42.4% for degradation), showing a significant correlation between the two phases (**Supp. Fig. 2I**). However, some compounds displayed opposite results in the two phases, possibly because in phase 2 we compiled the effect of three different concentrations. The compounds were categorized in 5 groups: pro-engulfment compounds:; pro-degradation, anti-engulfment, pro-degradation compounds; and a compound with mixed pro-engulfment and anti-degradative properties. Their chemical structure and main therapeutic use are shown in **Fig. 1D**. The mechanisms of action of these compounds were multiple, but several of them targeted main neurotransmitter pathways, such as dopamine, serotonin, adrenaline, noradrenaline, and GABA (trifluoperazine, nafronyl oxalate, maprotiline, dobutamine, baclofen). Another group of compounds targeted sodium and potassium channels (gliclazide, nicardipine, zoxazolamine). A final group of compounds targeted pathways related to DNA and RNA and are used as antitumoral agents (N6-methyladenosine, thioguanosine, doxorubicin).

**Figure 1.**
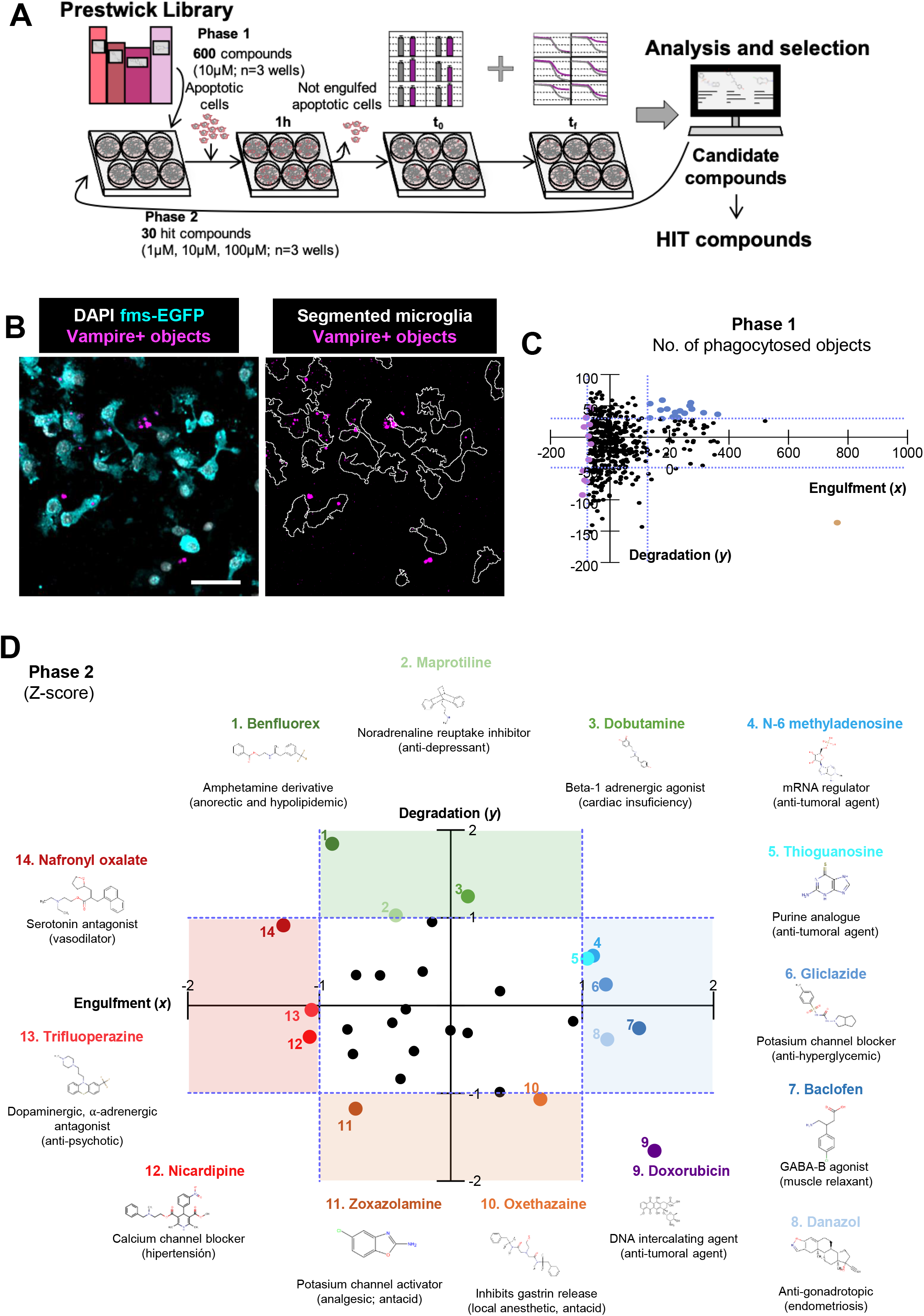
A-HTS screening and hit compounds. **A**) Cartoon depicting the experimental paradigm for A-HTS followed during this project. **B**) Representative fluorescence image from fms-EGFP microglia (lefts) and colocalization between Vampire+ objects and segmented microglia (right). Scale bar=50μm. **c**) Dot plot comparing the effects of the treatments on engulfment and degradation of apoptotic cells. In blue, compounds selected as candidate potentiators of phagocytosis. In mauve, compounds selected as candidate inhibitors of phagocytosis. In beige is doxorubicin which displays pro-engulfment and anti-degradation effects. Dotted lines represent mean number of phagocytosed objects per microglia ± standard deviation. **D**) Compounds that potentiate engulfment (blue) or degradation (green), and inhibit engulfment (red) or degradation (orange) of apoptotic cells. The results were deemed significant if their engulfment or degradation z-scores were higher or lower than 1 (dotted purple lines).

### Roche High Throughput screening (R-HTS)

In a first step, we developed an imaged based 384 well assay to measure the phagocytic activity of iPSC-derived macrophage-like cultures in the alpha-synuclein overexpressing SFC831-03-03 3xSNCA cell line. Pre-macrophages were differentiated to macrophages in the final assay format, incubated with compounds for 24 hrs and pH-Rodo labeled apoptotic NPCs for the last two hours of incubation. Macrophage-like nuclei were detected with Hoechst and the cytoplasm was detected by its reflection. Cells were classified as phagocytotic when they contained 1 or more pHrodo spots in the cytoplasmic region (**Fig. 2A**). Standard performance assays were assessed by blocking NPC phagocytosis with cytochalasin D and by inducing zymosan phagocytosis by opsonizing with fetal calf serum (FCS)(**Fig. 2B, C**). The phagocytic activity was reduced in macrophages derived from the 3xSNCA line compared to cells derived from an apparently healthy donor (SBNeo), confirming prior observations (Haenseler et al., 2017) (**Fig. 2D**), and supporting our choice to use the 3xSNCA line to find phagocytosis enhancers.

**Figure 2.**
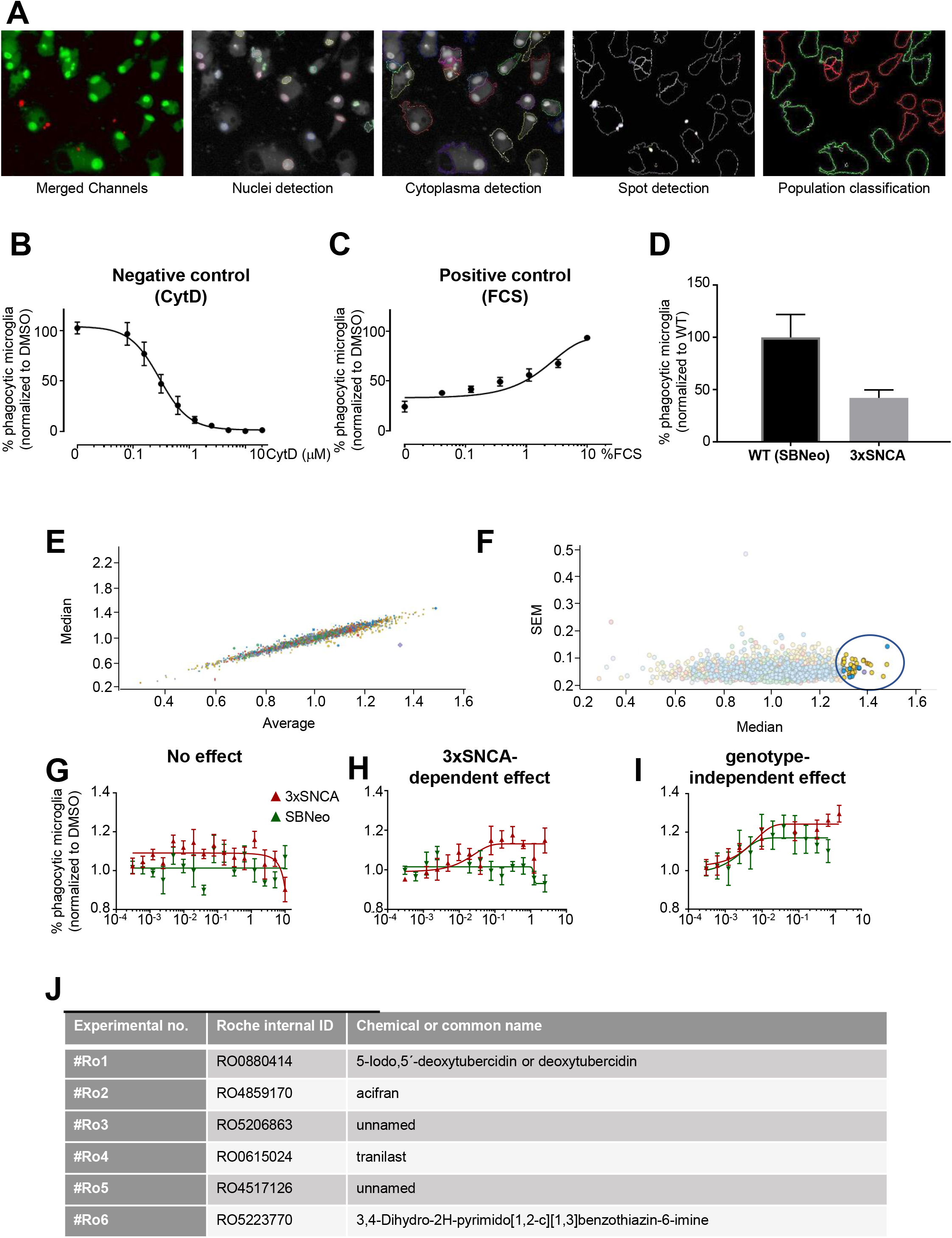
R-HTS screening, controls, and hit compound identification. **A)** Representative image of iPSC-derived macrophages phagocytosing pHrodo labeled NSC and the masks for detecting nuclei, cytoplasma, spots and categorization of phagocytic (green) and non phagocytic (red) cells. **B)** Inhibition of phagocytic activity of zymosan in macrophages pre-treated with cytochalasin D for 24h, normalized to DMSO treated control. **C)** Stimulation of phagocytosis by opsonisation of zymosan with FCS **D)** Phagocytosis in WT (SBNeo) and 3x-SNCA macrophages derived from different donors. Cells were incubated with 3 times the amount of fixed, pHrodo labeled NSCs. **E)** Comparison of the median vs the average effect on 6 replicates of the 2,200 compounds at 1μM on the phagocytosis of macrophages. Each dot represents one compound, the color indicates the compound source plate and the size of the symbol the standard deviation between replicates (n=6). **F)** Hit compounds from the primary screen were identified as those with phagocytosis above 1.3 relative to control and SEM below 0.2 (blue circle) from the 6 replicates. **G-I)** Examples for outcomes in a dose-response experiment(six 1:2 dilutions starting at 10μM) of primary hits comparing WT (SBNeo)- and 3xSNCA-derived microglia/macrophages showing no confirmation of phagocytosis modulation (**G**), an effect only in 3xSNCA cells (**H**), and a global phagocytosis modulatory effect in both SBNeo and 3xSNCA cells (**I**). **J)** Summary table of screening hits that were progressed to testing in organotypic slice cultures.

The primary screen was performed at 1 uM with 6 replicates on macrophage-like cells derived from the 3xSNCA line. The % of phagocytic microglia was normalized over the DMSO control in each plate and averaged over replicates. The comparison of Median vs Average showed a strong correlation with only few outliers (**Fig. 2E**), indicating a high reproducibility and robustness of the assay. The assay hits that were potential inducers of phagocytosis were identified as those compounds with a ratio of phagocytic microglia higher than 1.3 compared to control and SEM below 0.2 (**Fig. 2F**). To confirm the hits and test whether their modulation of phagocytosis was related to SNCA, a 16 point dose respone (0.0001 to 10 uM?) was run on macrophages generated from both SBNeo and 3xSNCA lines. We discriminated in three possible outcomes, no confirmation of phagocytosis modulation(**Fig. 2G**), selectively inducing phagocytosis in 3xSNCA (**Fig. 2H**) and general inducers of phagocytosis (**Fig.2I**). From this last group, 6 compounds were selected for further validation in organotypic cultures (**Fiig. 2 J**).

### Organotypic culture validation of A-HTS and R-HTS compounds

Next, we tested both the A-HTS and the R-HTS candidate compounds in hippocampal organotypic slices, which retain all the cell types from the CNS, as well as the structures and connectivity (Beccari et al., 2018). We incubated the organotypic slices with the compounds at 100μM (the highest dose used in the A-HTS) for 6 hours and analyzed cell death, microglial numbers, microglial phagocytosis of dead (apoptotic and necrotic) cells, and microglial morphology. We used nuclear morphology and propidium iodide (PI) to distinguish between apoptotic (pyknotic, PI^-^) from primary necrotic cells (not pyknotic, PI^+^), and secondary necrotic cells resulting from evolved apoptotic cells (pyknotic, PI^+^). As necrotic cells were very scarce, all PI+ cells were pooled together.

The candidate compounds had been categorized in the A-HTS into four different groups based on their effects on engulfment and degradation separately. However, analysis of phagocytosis in organotypic slices does not allow discriminating between the effects on engulfment or degradation. Instead, we determine the Ph index as a measure of the net effect (engulfment-degradation). Therefore, both pro-engulfment and anti-degradation compounds would increase the Ph index, and pro-degradation and anti-engulfment compounds would decrease it. These seemingly counterintuitive scenarios can be discriminated by taking into account the total number of dead cells, which should be expected to decrease in pro-engulfment and pro-degradation compounds, and increase in anti-engulfment and antidegradation compounds.

Due to technical reasons related to the vehicle in which the compounds were dissolved and the number of compounds, we divided the experiments into different batches. As controls, we used untreated (UT) slices and the vehicle in which the compounds were dissolved. First, we tested compounds dissolved in DMSO: potential phagocytosis enhancers Roche compounds #Ro1-6, thioguanosine, N6-methyladenosine, danazol, and gliclazide, potential phagocytosis inhibitors oxethazaine and zoxazolamine, and the mixed-effect compound doxorubicin. The enhancer maprotiline and the blocker nicardipine were dissolved in methanol, and the enhancer dobutamine and the blockers trifluoperazine and nafronyl were dissolved in water.

We first assessed Roche compounds, and we found that deoxytubercidin (#Ro1), a ribose derivative, highly increased the number of apoptotic and PI^+^ cells (**Fig. 3A-C**), and strongly inhibited their phagocytosis (**Fig. 3D-E**). No effect on the number of microglia was found (**Fig. 3F**), suggesting that deoxytubercidin did not have a global toxic effect but acted as a phagocytosis blocker, in contrast to its initial categorization as a phagocytosis enhancer in the R-HTS. Deoxytubercidin also increased the morphological complexity of microglia (**Fig. 3F-J**). In the second batch of Roche compounds, we did not observe any significant changes in cell death, phagocytosis, or microglia numbers compared to the DMSO vehicle but not to UT slices (**Supp**. **Fig. 3A-F**). We only found a small reduction in the phagocytosis of PI+ cells by Ro4 (**Supp**. **Fig. 3E**) and a reduction in microglial complexity by Ro6 compared to DMSO (**Supp**. **Fig. 3G-J**). Overall, these results identified deoxytubercidin as a phagocytosis inhibitor in organotypic slices.

**Figure 3.**
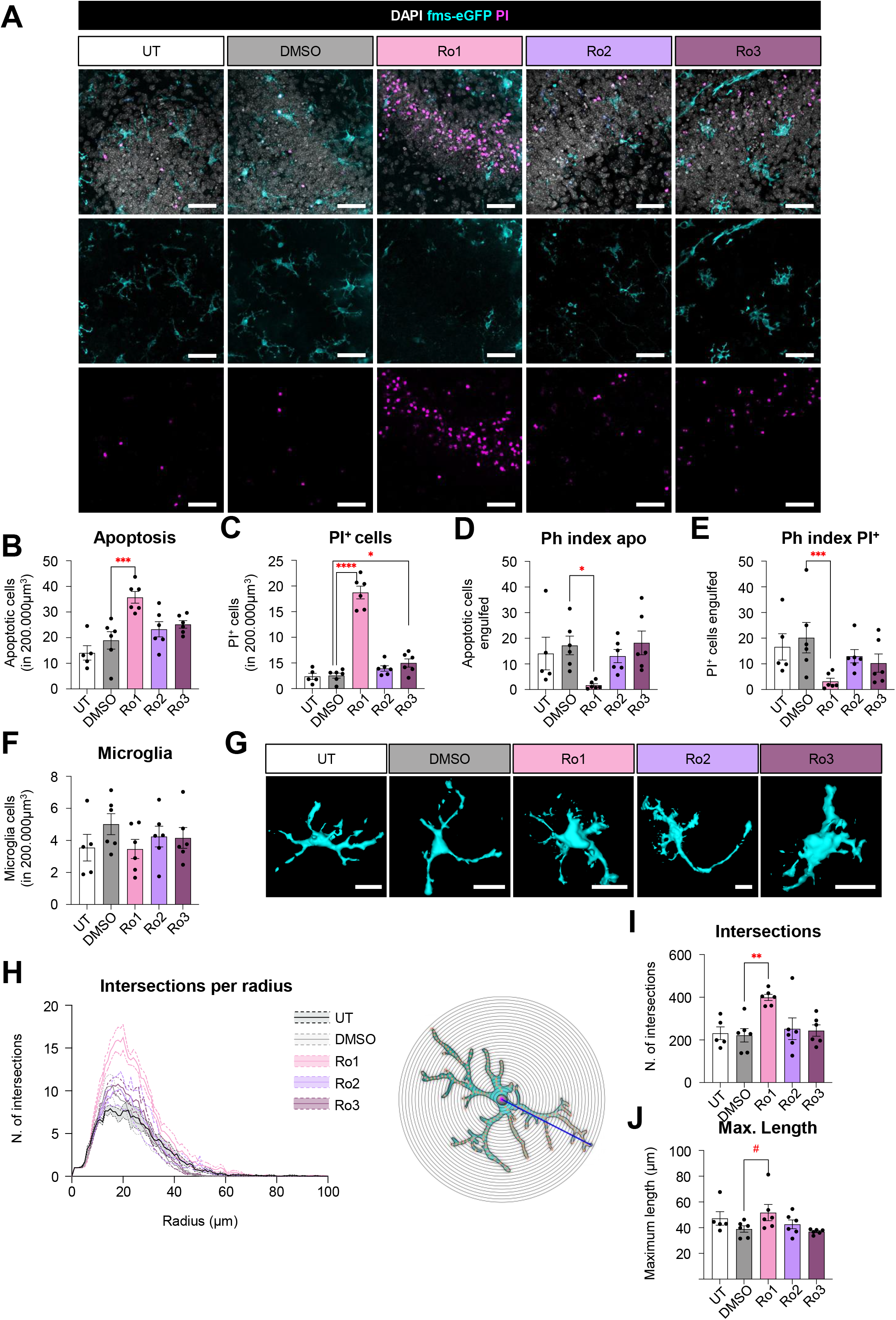
Ro1 (deoxytubercidin), but not Ro2 or Ro3, inhibits microglial phagocytosis in organotypic slices. **A**) Representative confocal z-stack of organotypic slices incubated for 6 hours with Ro1 (deoxytubercidin), Ro2, and Ro3, as well as Untreated (UT) and DMSO (1:1000% in cell culture medium), showing cell nuclei stained with DAPI (white), microglia (cyan), and PI^+^ cells (magenta). **B, C**) Number of apoptotic and PI^+^ cells in 200.000μm^3^, respectively. **D, E**) Percentage of apoptotic and PI^+^ cells engulfed by microglia (Ph index), respectively. **F**) Number of microglia cells in 200.000μm^3^. **G**) 3D reconstructions of representative morphological features of microglia for each tested compound. Note the variable degrees of dendrite branching, number and lengths of processes, and soma sizes for microglia exposed to different compounds. **H**) Histogram showing the number of microglial intersections in each radius starting from the soma using Sholl analysis. Concentric circles are drawn from the centre/soma of the cell (magenta dot). An intersection (orange dot) is considered whenever a concentric circle intersects a microglial process. Consequently, the total of all orange dots represents the sum of intersections. The radius of the last concentric circle intersecting a process is considered the maximum radius/length (blue line). **I**) Sum of all intersections of microglial processes. **J**) Maximum length of microglial processes. Scale bars in **A** and **G**: 50μm. In **A**, z=10.5μm. Bars show mean ± SEM. n=6 (UT), n=6 (DMSO), n=6 (Ro1), n=6 (Ro2), and n=6 (Ro3), where n is the number of animals. In **B**, **D**, **F**, and **I**, data were analyzed with a one-way ANOVA followed by a Dunnett’s multiple comparisons test (*vs* DMSO). In **C**, **E**, and **J**, data were SQRT, Log10, and Log10 transformed, respectively, to comply with normality and homoscedasticity and then analyzed with a one-way ANOVA followed by a Dunnett’s multiple comparisons test (*vs* DMSO). * represents p < 0.05; ** represents p < 0.01; *** represents p < 0.001; **** represents p < 0.0001.

In the next batch, we assessed two potential engulfment enhancers that shared a mechanism of action with deoxytubercidine: thioguanosine, a purine analogue used as an antineoplastic compound (De, 2023); and N6-methyladenosine, an mRNA regulator involved in tumorigenesis and compound resistance (He et al., 2019; Wang et al., 2023). Both compounds reduced apoptotic cells phagocytosis but only m6A increased the number of dead cells or microglia (**Supp**. **Fig. 4A-E**), without changes in microglial number of morphology (**Supp**. **Fig. 4F-J**). As deoxytubercidin, both thioguanosine and N6-methyladenosine behaved as phagocytosis inhibitors in organotypic slices, although they did not lead to delayed apoptotic cell clearance and accumulation, possibly because the effects on phagocytosis were stronger in deoxytubercidin. We then analyzed a compound with mixed pro-engulfment and anti-degradation properties in the A-HTS, doxorubicin, an anthracycline antibiotic used as an antineoplastic agent (Carvalho et al., 2009; Johnson-Arbor and Dubey, 2023). As doxorubicin presents an intrinsic autofluorescence (Shah et al., 2017), which overlaps with the signal of PI and prevents DAPI staining, we used its fluorescence as a marker of cell nuclei. Doxorubicin showed a non-significant trend (p=0.2276 compared to DMSO) towards decreased number of apoptotic cells and decreased phagocytosis (**Supp**. **Fig. 5A-C**), which could be related to the reduced microglial morphological complexity (**Supp**. **Fig. 5E-H**). The effect on phagocytosis can be interpreted as a reduction in engulfment or an increased degradation. Overall, the three compounds in this batch, and thioguanosine, N6-methyladenosine, and doxorubicin behaved as phagocytosis inhibitors in organotypic slices similar to deoxytubercidine, regardless of their initial classification in the A-HTS. From this initial set of experiments, we analyzed the effect of DMSO compared with UT cultures. Because no global effects of DMSO across experiments were observed in the Ph index (13.8 ± 1.9 vs 18.2 ± 1.2, p=0.09, with no statistically significant effect in each individual experiment), we skipped it in the rest of the compounds dissolved in DMSO (danazol, gliclazide, oxethazaine, and zoxazolamine).

In the next batch, we analyzed two compounds that were classified as pro-engulfment compounds in the A-HTS: danazol, an anti-gonadotropic and anti-estrogenic agent (Barbieri and Ryan, 1981); and Gliclazide, an anti-hyperglycemic compound (Khunti et al., 2020). However, danazol did not exert significant effects on cell death, phagocytosis, microglia number or morphology, and gliclazide only increased apoptosis and showed a trend towards a decreased phagocytosis (**Supp**. **Fig. 6**). We then analyzed two compounds classified in the A-HTS as degradation blockers: oxethazaine, a local anesthetic and antacid used to relief pain associated with the digestive system (Yasuhara and Levy, 1988; McMillen et al., 1992)(Seifter et al., 1962), and zoxazolamine, a centrally-acting muscle relaxant (Yasuhara and Levy, 1988; McMillen et al., 1992). Oxethazaine increased cell death but did not affect apoptotic cell phagocytosis, and had no effect on microglia number or morphology, suggesting a toxic effect unrelated to phagocytosis (**Supp**. **Fig. 7**). Zoxazolamine did not alter the number of apoptotic cells (PI was not added in this particular experiment) but increased the Ph index (**Supp**. **Fig. 8**), an effect that can be interpreted as a reduction in degradation (in agreement with its initial classification as a degradation blocker in the A-HTS), or as an increase in engulfment. Zoxazolamine did not change the number of microglia but increased their morphological complexity (**Supp**. **Fig. 8E-H**), which could be related to increased sampling capacity and increased engulfment. From these experiments, we concluded that danazol and gliclazide had no relevant effect on phagocytosis, oxethazaine was toxic, and zoxaxolamine had a potential pro-phagocytosis effect, but its hepatotoxic effects (Eisenstadt and Elster, 1961) discouraged us from its further analysis *in vivo*.

Next, we assessed two compounds dissolved in methanol, maprotiline and nicardipine, that were originally classified as pro-degradation and anti-engulfment in the A-HTS, respectively. Maprotiline is a norepinephrine reuptake inhibitor used as an antidepressant (Wells and Gelenberg, 1981). This compound increased the number of dead cells and reduced their phagocytosis, without changing microglial numbers but reducing their complexity (**Supp**. **Fig. 9**). Because we found no significant effect of methanol compared to UT cultures, we skipped this group in the analysis of nicardipine, a calcium channel blocker used to treat hypertension and angina (Sorkin and Clissold, 1987). Similar to maprotiline, nicardipine increased the number of apoptotic cells and decreased their phagocytosis, without compromising microglial numbers or complexity (**Supp**. **Fig. 10**). These results suggest that both maprotiline and nicardipine were phagocytosis inhibitors in organotypic slices.

Finally, we analyzed compounds dissolved in water, the potential degradation enhancer dobutamine, and the engulfment blockers trifluoperazine and nafronyl, as initially classified in the A-HTS. Dobutamine, a beta-1 agonist used to treat heart diseases (Tuttle and Mills, 1975; Ashkar et al., 2023), increased the number of dead cells but only affected the phagocytosis of PI+ cells, and did not change microglial numbers but reduced their complexity (**Supp**. **Fig. 11**), suggesting a toxic effect. Trifluoperazine, a dopaminergic and α-adrenergic antagonist (Huerta-Bahena et al., 1983) used as an antipsychotic and anxiolytic (Howland, 2016); and nafronyl, a vasodilator and serotonin antagonist used in the management of vascular disorders (Aly et al., 2009) increased cell death, and trifluoperazine blocked the phagocytosis of PI+ cells and showed a trend to inhibit the phagocytosis of apoptotic cells (**Fig. 4A-E**). Moreover, both trifluoperazine and nafronyl reduced the complexity of microglia cells without changing their numbers (**Fig. 4F-J**). These findings indicated that nafronyl had a toxic effect, whereas trifluoperazine behaved as an anti-engulfment compound in organotypic slices.

**Figure 4.**
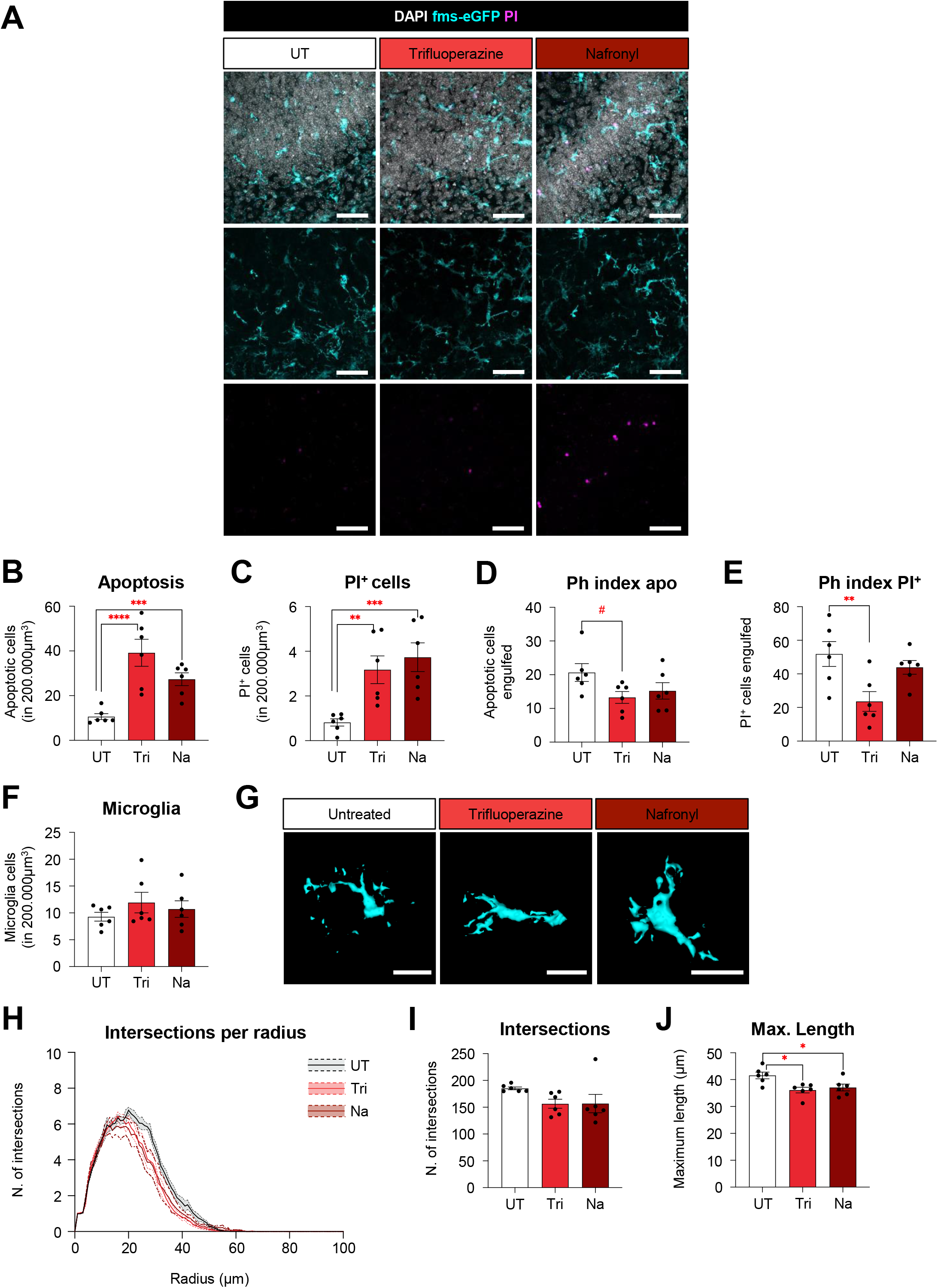
Trifluoperazine, but not nafronyl, inhibits microglial phagocytosis in organotypic cultures. **A**) Representative confocal z-stack of organotypic slices treated with trifluoperazine (Tri) and nafronyl (Na), as well as Untreated (UT) conditions, showing cell nuclei stained with DAPI (white), microglia (cyan), and PI^+^ cells (magenta). **B, C**) Number of apoptotic and PI^+^ cells in 200.000μm^3^, respectively. **D, E**) Percentage of apoptotic and PI^+^ cells engulfed by microglia (Ph index), respectively. **F**) Number of microglia cells in 200.000μm^3^. **G**) 3D reconstructions of representative microglia for each condition. **H**) Histogram showing the number of microglial intersections in each radius starting from the soma. **I**) Sum of all intersections of microglial processes. **J**) Maximum length of microglial processes. Scale bars in **A** and **G**: 50μm. In **A**, z=10.5μm. Bars show mean ± SEM. n=6 (UT), n=6 (Tri), and n=6 (Na), where n is the number of animals. In **D-F** and **I-J**, data were analyzed with a one-way ANOVA followed by a Dunnett’s multiple comparisons test (vs control). In **B-C**, data were Log10 and SQRT transformed, respectively, to comply with normality and homoscedasticity and then analyzed with a one-way ANOVA followed by a Dunnett’s multiple comparisons test (*vs* Untreated). # represents p < 0.1; * represents p < 0.05; ** represents p < 0.01; *** represents p < 0.001; **** represents p < 0.0001.

**Figure 5.**
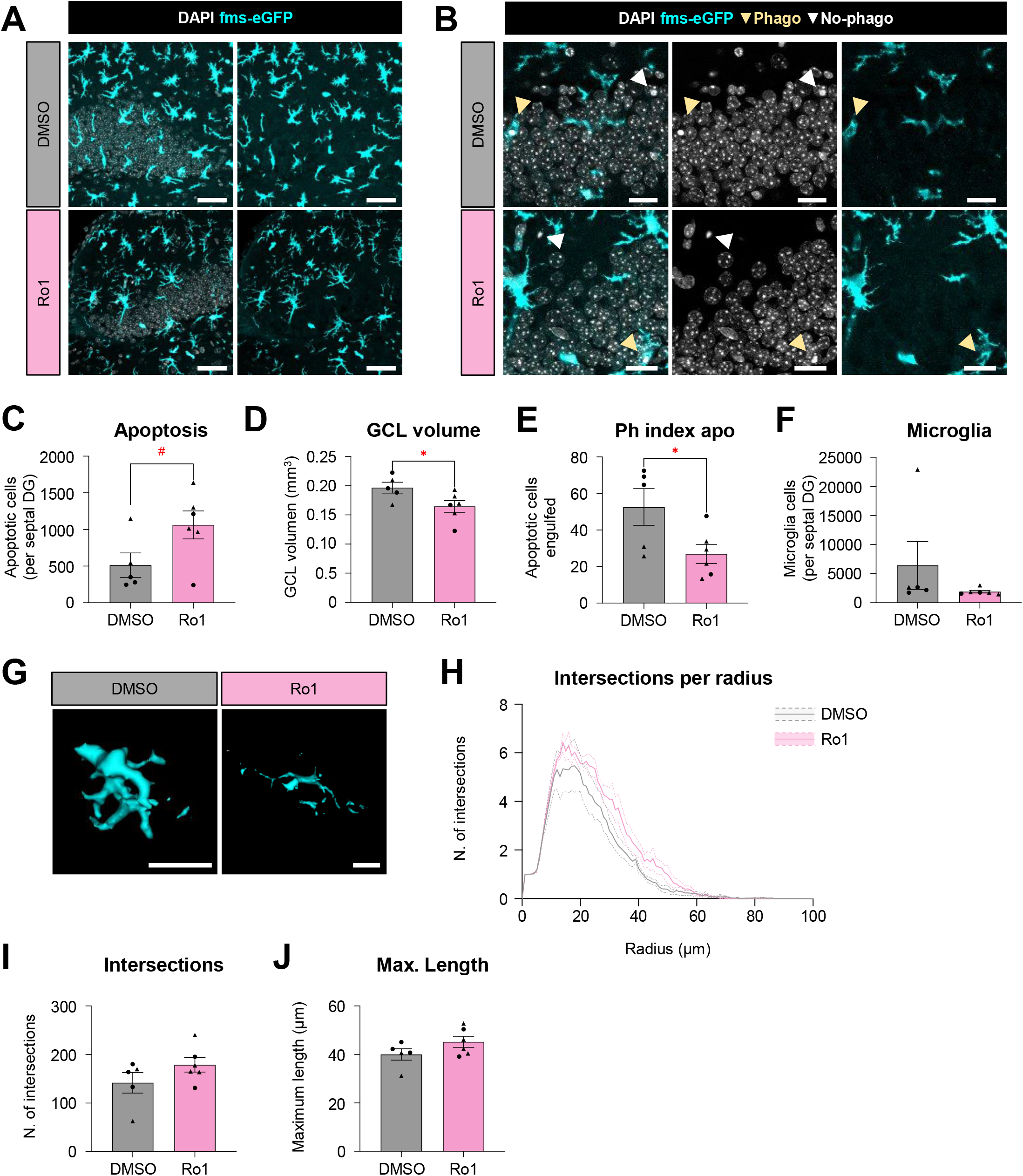
Ro1 (deoxytubercidin) inhibits microglial phagocytosis *in vivo*. **A**) Representative confocal z-stack of the hippocampus of mice administered with Ro1 and its vehicle DMSO showing cell nuclei stained with DAPI (white) and microglia (cyan). **B)** Confocal z-stacks showing examples of apoptotic cells phagocytosed (yellow arrowheads) and non-phagocytosed (white arrowheads) by microglia (cyan) in each condition. Apoptotic cells were identified by their condensed chromatin (pyknosis) using DAPI staining (white). **C**) Number of apoptotic cells in the septal GCL of the hippocampus. **D)** Volume of the septal GCL in each condition. **E**) Percentage of apoptotic cells engulfed by microglia (Ph index). **F**) Number of microglia cells in the septal GCL. **G**) 3D reconstructions of microglia. **H**) Histogram showing the number of intersections in each radius starting from the soma. **I**) Sum of all intersections of microglial processes. **J**) Maximum length of microglial processes. Scale bars in **A** and **G**: 50μm; in **B**: 20μm. In **A**, z=10.5μm; in **B**: z=4.2μm (DMSO) and z=5.6μm (Ro1); Bars show mean ± SEM. n=5 (DMSO) and n=6 (Ro1). In **C-E** and **I-J**, statistical significance was analyzed with an unpaired t-test. In **F**, data were *sin* transformed to comply with normality and homoscedasticity and then statistical significance was analyzed with an unpaired t-test. # represents p=0.06; * represents p < 0.05.

**Figure 6.**
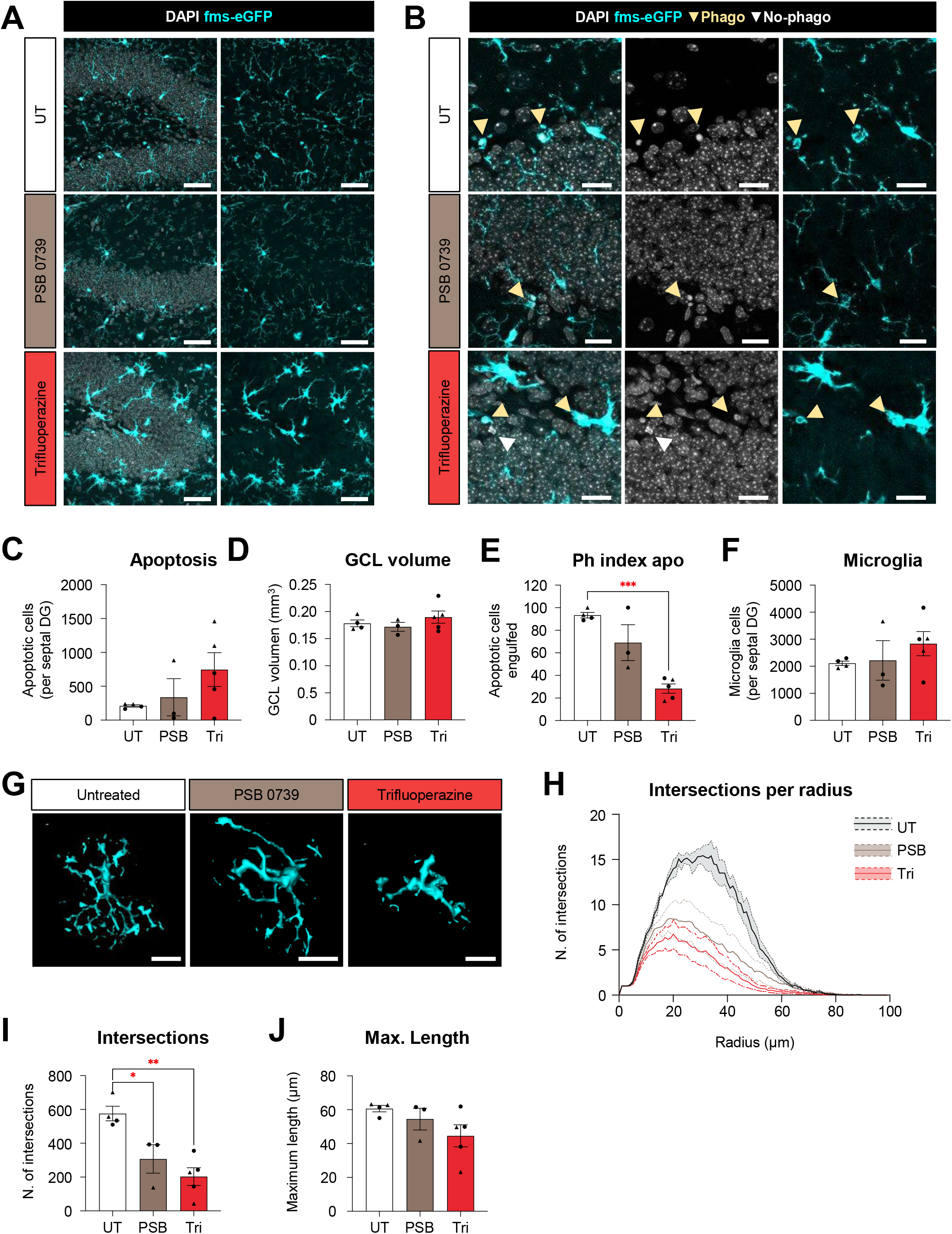
Trifluoperazine inhibits microglial phagocytosis *in vivo* more robustly than PSB-0739. **A**) Representative confocal z-stack of the hippocampus of mice administered with trifluoperazine (tri), PSB-0739 (PSB), and its vehicle DMSO showing cell nuclei stained with DAPI (white) and microglia (cyan). **B)** Confocal z-stacks showing examples of apoptotic cells phagocytosed (yellow arrowheads) and non-phagocytosed (white arrowheads) by microglia (cyan) in each condition. Apoptotic cells were identified by their condensed chromatin (pyknosis) using DAPI staining (white). **C**) Number of apoptotic cells in the septal GCL of the hippocampus. **D)** Volume of the septal GCL in each condition. **E**) Percentage of apoptotic cells engulfed by microglia (Ph index). **F**) Number of microglia cells in the septal GCL. **G**) 3D reconstructions of microglia. **H**) Histogram showing the number of intersections in each radius starting from the soma. **I**) Sum of all intersections of microglial processes. **J**) Maximum length of microglial processes. Scale bars in **A** and **G**: 50μm; in **B**: 20μm. In **A**, z=10.5μm; in **B**: z=7μm (UT); z=7μm (PSB); and z=4.2μm (tri). Bars show mean ± SEM. n=4 (UT), n=3 (PSB), and n=5 (tri). In **C-J**, data were analyzed with a one-way ANOVA followed by a Tukey’s post hoc test. * represents p < 0.05; ** represents p < 0.01; *** represents p < 0.001.

**Figure 7.**
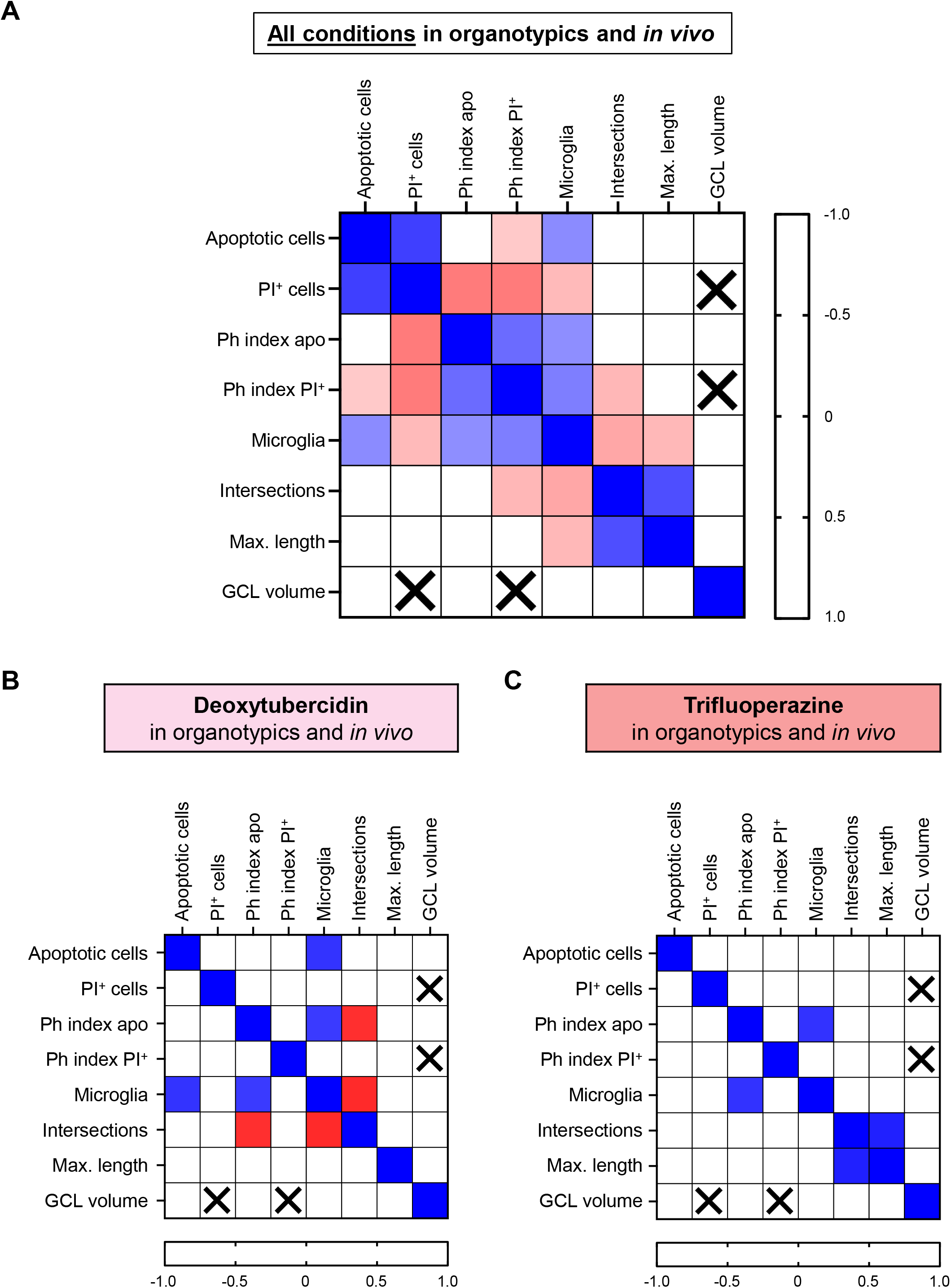
Multiple Correlation analysis does not reveal evident relations between microglial phagocytosis and morphology across all experiments, neither upon treatment with deoxytubercidin nor trifluoperazine. **A)** Correlation matrix analysis across all experiments including all conditions (untreated and treated conditions) in organotypic slices and *in vivo*. **B**) Correlation matrix analysis of organotypic slices and animals treated with deoxytubercidin. **C**) Correlation matrix analysis of organotypic slices and animals treated with trifluoperazine. In **A**, **B**, and **C:** cells with an X represent correlations that were not carried out. Only correlations with p < 0.01 are indicated, to reduce possible random detection of significant correlations due to multiple comparisons. Colored squares represent the Spearman’s correlation coefficient ρ, represented in a scale ranking from ρ = -1.0 to 1.0.

In summary, hippocampal organotypic slices allowed us to validate some compounds identified in the A-HTS and R-HTS in a complex *ex vivo* system that maintains neuronal connectivity and microglia function. Most compounds (deoxytubercidin, thioguanosine, N6-methyladenosine, doxorubicin, gliclazide, maprotiline, nicardipine, nafronyl, trifluoperazine), regardless of their original classification in previous HTS, behaved as anti-engulfment compounds: they decreased the Phagocytic index and increased the number of dead cells. Thus, these results strongly suggest that it is easier to inhibit than to enhance microglial phagocytosis. The only compounds with consistent effects between the HTS and the organotypic cultures were nicardipine and trifluoperazine, both confirmed anti-engulfment compounds.

### *In vivo* validation of A-HTS and R-HTS compounds

Following our bottom-up strategy in the validation of microglial phagocytosis modulators, we tested *in vivo* the compounds with the most robust effects on the inhibition of engulfment: deoxytubercidin and trifluoperazine. As a model, we used the adult hippocampal neurogenic niche, where newborn cells undergo apoptosis and are phagocytosed by microglia in physiological conditions (Sierra et al., 2010). We tested deoxytubercidin, trifluoperazine, and the P2Y12R antagonist PSB-0739 (Cserép et al., 2020) as a positive control for phagocytosis inhibition. Compounds were administered intrahippocampally for 2 days at 10μM concentration using osmotic minipumps (injected volume: 100μl) to 2-month-old mice and analyzed cell death, microglia number, phagocytosis of apoptotic cells, and microglial morphology in the granule cell layer (GCL) of the hippocampus. As control groups, we used untreated (UT) mice for trifluoperazine and PSB-01739, and DMSO-treated mice for Deoxytubercidin.

Deoxytubercidin showed a non-significant tendency towards increased apoptosis compared to the DMSO control, which was probably related to a reduced GCL volume that could suggest a potential toxic effect. Deoxytubercidin also showed a signficant 48.7 ± 9.9% reduction in the phagocytosis efficiency determined by the Ph index, which was not accompanied by significant alterations in microglial numbers of morphology (**Fig. 8**). Trifluoperazine, on the other hand, had no global effects on apoptosis nor GCL volume, but showed a more robust reduction in phagocytosis than PSB-0739 (69.7 ± 4.4% vs 26.0 ± 23.1% reduction). Both trifluoperazine and PSB-0739 effectively reduced microglial complexity without affecting their numbers (**Fig. 9**). Together, these results demonstrate that both deoxytubercidin and trifluoperazine can efficiently inhibit microglial phagocytosis in this adult hippocampal neurogenic niche apoptosis *in vivo* paradigm.

Finally, we performed a multiple correlations analysis to explore whether changes in microglial numbers and morphology could be responsible for microglial phagocytosis inhibition. Overall, across all compounds and experiments (in organotypics and in vivo), we found a significant correlation of the Ph index with microglial numbers (**Fig. 10A**). For all deoxytubercidin experiments combined, we did find an additional significant negative correlation between the Ph index of apoptotic cells and the number of microglial intersections (**Fig. 10B**), suggesting that the phagocytosis impairment induced by deoxytubercidin could be related to reduced morphological complexity. However, we did not find any correlation between microglial phagocytosis and morphology in all the experiments with trifluoperazine combined (**Fig. 10C**). Therefore, our analyses indicate that deoxytubercidin and trifluoperazine may affect microglial phagocytosis through different mechanisms, as their known mode-of-action would suggest (i.e., purine nucleoside analoge vs dopaminergic antagonist, respectively).

## DISCUSSION

Glial cells, including microglia, have been long recognized as therapeutic targets for neurodevelopmental, neurological, and neurodegenerative disorders (Paolicelli et al., 2022). Microglia are key players in health and disease by controlling and executing the inflammatory response and phagocytosing different types of cargo, from apoptotic cells and unwanted spines to myelin debris and protein aggregates. While there are multiple immune modulators, both approved and in clinical trials, microglial phagocytosis has an unexplored therapeutic potential in conditions where it is dysfunctional, such as epilepsy or stroke (Abiega et al., 2016; Sierra□Torre et al., 2020; Beccari et al., 2023), or when it is exacerbated, such as in ischemia or after exposure to certain amyloid-beta species (Neher et al., 2011, 2013; Neniskyte and Brown, 2013). To explore how phagocytosis in microglia could be addressed pharmacologically, we have employed an unbiased bottom-up approach staring with a phenotypic (i.e., phagocytosis) small molecule HTS in mouse primary microglia (A-HTS) and human iPSC-derived macrophages (R-HTS), followed by initial testing of a few hits in an organotypic hippocampal slice cultures paradigm of apoptotic cell phagocytosis, and confirmation of an effect for two compounds of interest in an in vivo paradigm of removal of apoptotic newborn cells in the adult hippocampal neurogenic niche in mice.

For the two HTS, we chose two separate chemical libraries which contain small molecules that are well-annotated related to their known mechanism. The choice of primary murine microglia and human iPSC-derived macrophages as a functional surrogate for microglia allowed for a translational comparison of outcomes between a model species and the disease-relevant human species. Our hit rate is within the reported range in a metanalysis of 400 HTS, ranging between 0.01% and 0.14% (Zhu et al., 2013). Overall, our approach exemplifies that well-annotated chemical libraries can be used to discover novel modes of action for small molecules with known mechanism. Here, we identified the antipsychotic/antineoplasic trifluoperazine and the ribose derivative deoxytubercidin as inhibitors of microglial phagocytosis in our experimental setup. These molecules and their mechanism could now be employed to explore further the pathways to modulate phagocytosis as an starting point for new drug discovery projects towards a pharmacological modulation of microglia in disease settings.

### Novel screening platforms for microglial functions

In the last few years, there has been a strong interest in developing platforms for testing microglial functions, using as models cell lines, such as the mouse BV2 (Patil et al., 2021), primary mouse cultures (Mason et al., 2023), human microglial cells derived from autopsy tissue (Smith et al., 2013), and microglia-like cells derived from circulating monocytes (Connor et al., 2022; Zhang et al., 2022) or from induced pluripotent stem cells (iPSCs) (Bassil et al., 2021). These platforms have focused on microglial functions, including the expression or release of inflammatory, immunomodulatory and/or neuroprotective factors (Hansen et al., 2010; Patil et al., 2021) or exovesicles (Ruan et al., 2022), or on the response to pathological protein aggregates derived from amyloid beta or alpha-synuclein (Bassil et al., 2021). Recent reports have also described high content screening platforms for pharmacological (Connor et al., 2022) or CRISPR-identified genetic (Chang et al., 2023) modulators of microglial phagocytosis and clearance of amyloid beta.

Our HTS specifically focused on phagocytosis of apoptotic cells. Based on our published protocol (Beccari et al., 2018), in A-HTS we used primary cultures and a fluorescently-labeled apoptotic neuronal cell line as substrate. Apoptosis was induced using staurosporine, and a high content imaging platform was employed to image phagocytosis at two time points, to allow discriminating between engulfment and degradation. In R-HTS, we used macrophages derived from an iPSC line donated by a person with an alpha-synuclein gene triplication, which causes a familial form of Parkinson’s disease. We confirmed in the Roche laboratory and assays setup that these 3xSNCA iPSC-derive macrophages indeed have an intrinsic deficiency in phagocytosis as previously reported (Haenseler et al., 2017). For the R-HTS we chose apoptotic neuronal debris labeled with pH-rodo, which only emits fluorescence in acidic environments thus confirming that the engulfed and internalized substrate is transported to the lysosome. In addition to the proper screening, here we also describe a complete analysis pipeline, including filtering steps for sources of variability (such as technical replicates and toxic effects); the use of a z-score to compile data from different concentrations; and validation of selected hit compounds in organotypic mouse hippocampal slice cultures and an *in vivo* paradigm of phagocytic removal of new born apoptotic cells in the neurogenic niche. We anticipate that this integrated pipeline can be useful for other researchers planning ahead their own HTS.

Several considerations must be taken into account when designing an in vitro HTS: 1, Substrate: we recommend using biologically relevant substrates (apoptotic cells, myelin debris, bacteria, etc, depending on the pathophysiological context studied), instead of using latex beads. When assessing apoptotic cell engulfment, pHrodo is helpful to assess that the substrate has reached the lysosome. 2, Timing: discriminating between engulfment *vs* degradation along a time course can provide additional information about the mechanism of action of the hit compounds. 3, HTS design: we recommend including reference compounds that induce or inhibit phagocytosis (i.e., a clear and best-defined regulator of function) or have no effect (inert, like a buffer control), commonly referred to as positive and negative controls. 4, Range of concentrations: researchers may opt to select hits that are more consistent across concentrations using a z-score (as in A-HTS) or those that show a dose-response effect (as in R-HTS). It is important to note that some compounds may only show a specific mode of action at low concentrations and hit off-targets or become toxic at higher concentrations. 5, Library diversity: the polypharmacology of the compounds and their known must be considered when selecting the chemical library. 6, Compound cell specificity: because neurons and microglia communicate bi-directionally, some compounds may indirectly regulate microglial phagocytosis by acting on neurons, which in turn may modulate microglial function. 7, Validation: it is imperative to validate the hit compounds in complex systems, specially *in vivo* or in 3D multicell-type human model systems. Relevant models of human pathologies must be used to accelerate the translatability of the findings.

### Trifluoperazine as microglial phagocytosis inhibitor

Our bottom-up approach and experimental setup allowed us to identify trifluoperazine as a phagocytosis inhibitor with stronger inhibitory effect at 100 µM than PSB-0739, a highly specific P2Y12R inhibitor (Cserép et al., 2020). Trifluoperazine is a phenothiazine, a first-generation heterocyclic antipsychotic compound that has recently been shown to display antitumoral effects as well (Kidron and Nguyen, 2024). It was also recently identified as a potential phagocytosis inhibitor in a machine learning-driven screening based on its *in silico* inhibitory activity of MEGF10 (Multiple EGF Like Domains 10) (Gravina et al., 2022), an astrocyte-enriched receptor of the complement C1q protein that mediates astrocytic synaptic phagocytosis (Chung et al., 2013). Trifluoperazine is largely thought to act by antagonizing dopaminergic and α1 adrenergic receptors, although it also interferes with Ca^2+^ binding proteins, such as calmodulin and calsequestrin, and intracellular Ca^2+^ channels such as the ryanodine receptors (Qin et al., 2009; Kidron and Nguyen, 2024). Here, we have not explored the potential mechanism of action of trifluoperazine on microglia, but it is unlikely mediated by dopaminergic, α1 adrenergic, or ryanodine receptor genes (Drd1-2, Adra1, and Ryr1-3), which are expressed at very low levels in microglia (https://brainrnaseq.org/). While trifluoperazine Ca^2+^ related targets may be responsible for the altered phagocytosis efficiency, this effect seems to be independent of an effect in microglial morphological complexity.

Trifluoperazine, commercialized as Stelazine, has been used to treat schizophrenia since the 1960s, and the consensus is that it reduces the activity of dopaminergic receptors in mesolimbic and mesocortical connections (Marques et al., 2004; Koch et al., 2014), although it may also act by regulating synaptic pruning by microglia or astrocytes (Germann et al., 2021). Trifluoperazine has several side effects, including alterations in striatal circuits that lead to pseudo-Parkinsonism and dystonia (Marques et al., 2004; Koch et al., 2014). Trifluoperazine also has anti-proliferative properties *in vitro* related to its ability to block calmodulin (Frankfurt et al., 1995), and recent studies have shown potent antineoplastic effects *in vivo* in several cancers, including glioblastoma (Kang et al., 2017). Our results, however, invite to speculate that the beneficial effect of trifluoperazine in cancer may be compromised if the resident phagocytes have impaired their ability to clean up apoptotic tumor cells because of the anti-phagocytic effect of the compound. Like many first generation antipsychotics, trifluoperazine seems to show clear signs of polypharmacology and off-target effects. Thus it is difficult to make any assumptions about the mode of action on different cell types and in different therapeutic settings and dosings. Still, trifluoperazine could be a valuable tool to study pathways that regulate phagocytosis, possibly opening new directions for identifying novel drug targets towards improving this important microglial function.

### RNA/DNA metabolism and microglial phagocytosis

Our screening also validated deoxytubercidin as a phagocytosis inhibitor in organotypic slices and *in vivo*, although it had been initially classified as enhancer in the R-HTS. Interestingly, the purine analog thioguanosine, and the mRNA regulator N6-methyladenosine were also initially classified as phagocytosis enhancers in A-HTS but were confirmed as inhibitors in organotypic slices. The mechanism and mode of action of these cytostatic and cytotoxic nucleoside analogues in microglia is unknown. Interestingly, recent data suggests a role of RNA- and DNA-binding proteins on microglial phagocytosis. For example, the RNA-binding protein Quaking (Qki), involved in RNA homeostasis, regulates myelin phagocytosis during cuprizone-induced demyelination (Ren et al., 2021). Similarly, IMP1 (IGF2 mRNA-binding protein, also known as ZBP1) regulates microglial phagocytosis of apoptotic cells during inflammation through binding to *actin* mRNA and regulating local protein translation (Imaz-Iruretagoyena et al., 2024). The TAR DNA-Binding Protein 43 (TDP-43) is involved in phagocytosis of amyloid plaques and dendritic spine debris in mouse models of amyloid pathology (Paolicelli et al., 2017). Future studies may want to explore how nucleoside analogues could lead to RNA or DNA damage, which in turn could change how regulatory proteins interact with phagocytosis regulating gene elements in microglia. It could then also be studied whether the differences observed in cell culture versus *in vivo* may come from the presence of several cell types that are simultaneously affected by the nucleoside analogues and may have changed their interaction due to RNA and DNA damage.

In summary, we demonstrate that based on well-annotated chemical librariers one can establish phenotypic screens to identify compounds with the ability to modulate the phagocytic function of microglia in vitro and in vivo. Our approach also shows the limitation of direct translation between effects observed in cell culture versus the ones in vivo. One reason why we detected mostly inhibitors of phagocytosis rather than enhancers could be that the cells in culture are already extremely efficient in that function and further optimization is difficult to achieve, even for iPSC-derived macrophages that show a clear deficit due to a genetic predisposition (i.e., 3xSNCA). Thus, for finding phagocytosis enhancers one may require *in vitro* platforms and novel human model systems mimicking specific pathological conditions that disrupt phagocytosis efficiency, such as oxygen and nutrient deprivation (for stroke), seizures (for epilepsy), of alpha-synuclein pathology and aggregates (for Parkinsońs disease). Microglial dysfunction in these different conditions is further influenced by additional aspects that are currently difficult to model, such as genetic predisposition and brain development, acute and strong inflammation, trauma-like events involving rupture or leakiness of brain vasculature, or a slow chronic degenerative progression with contriubtion of aging. The compounds identified in our screens are a starting point to study microglia function which may be regulated by thusfar unknown molecular mechanisms. Exploring the pathways affected by these molecular mechanisms in simple or next generation human model systems could help unravel the regulatory mechanisms behind phagocytosis in microglia with the goal to identify new pharmacological approaches to modulate microglia in disease.

## Data availability

Data is available upon request to

## Funding acknowledgements

This work was supported by grants from the Spanish Ministry of Science and Innovation Competitiveness MCIN/AEI/ 10.13039/501100011033 and by “ERDF A way of making Europe” (RTI2018-099267-B-I00 and PID2022-136698OB-I00), Basque Government grants (PIBA 2020_1_0030 and IT1473-22), and a BBVA Foundation Leonardo Award (IN[16]_BBM_BAS_0260) to AS. SG was initially supported by a Roche Postdoctoral Fellowship (RPF). IP performed some experiments at Roche as student intern under the Roche Internships of Scientific Exchange (RiSE) program. NRI was supported by a fellowship of the University of the Basque Country EHU/UPV.

