## Supplementary Text for "A bottom-up approach identifies the antipsychotic and antineoplastic trifluoperazine and the ribose derivative deoxytubercidin as novel microglial phagocytosis inhibitors"

**SUPPLEMENTARY FIGURES AND LEGENDS**

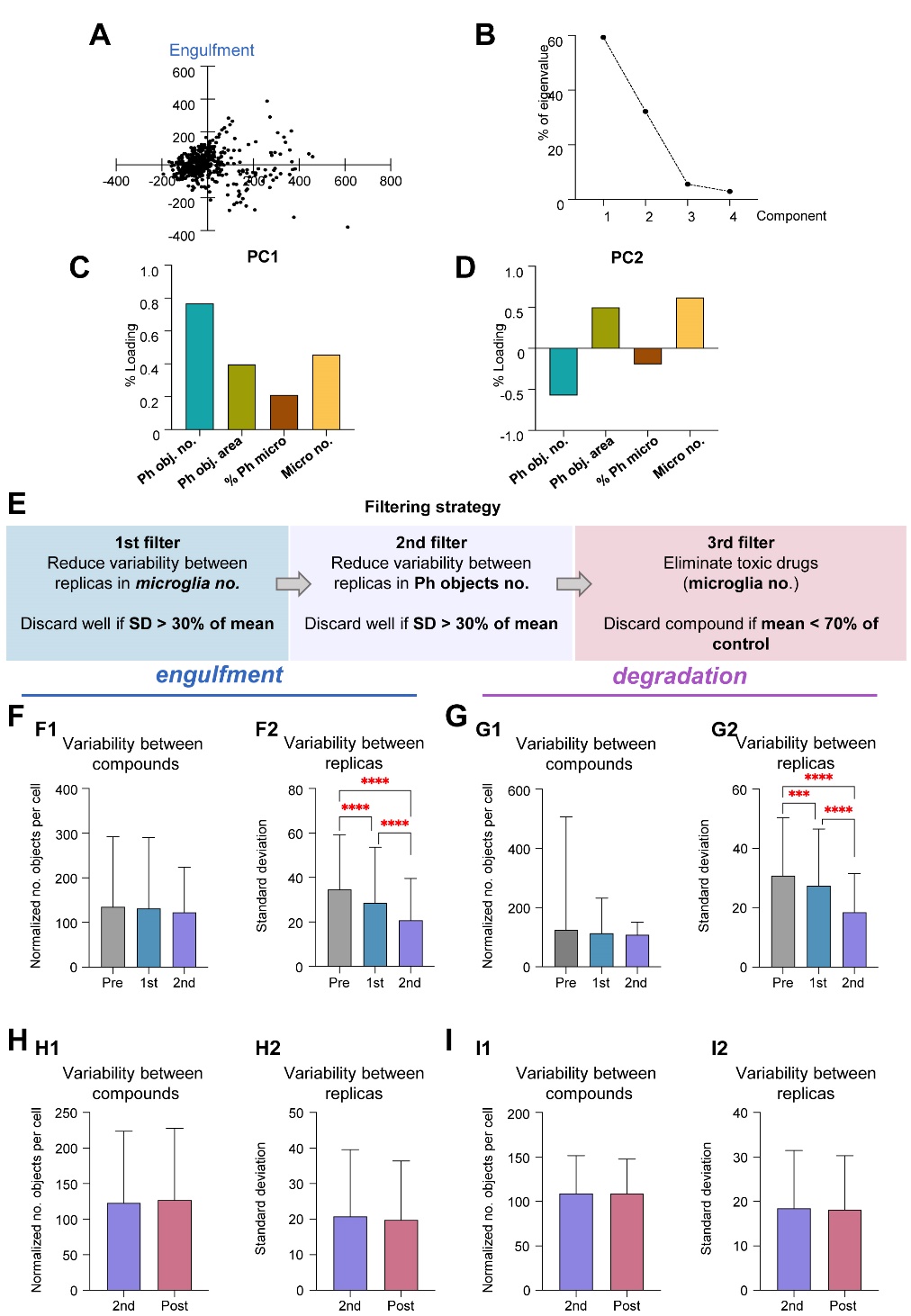

**Supplementary Figure 1. A-HTS phase 1 screening strategy.** **A**) PC1 vs PC2 dot plot representing all 600 compounds tested at 10μM. **B**) Graphical representation of the percentage that each PC represents on the total variance. Eigenvalues are the magnitude (the variance) associated with each PC. **C, D**) Bar graph representing the weight that each variable has on PC1 (G) or PC2 (H): number of phagocytosed objects (Ph obj. no.), phagocytosed objects area (Ph obj. area), % of phagocytic microglia (% Ph micro) and number of microglia (Micro no.) Loadings represent the correlations between variables and principal components, providing insights into the magnitude and direction of each variable's contribution to a determined component. The variable with the highest loading percentage on PC1 was the number of phagocytosed objects (Ph obj. no.), and was thus selected for the downstream analysis. **E**) Filtering strategy followed to reduce variability. **F, G**) Comparison of variability between compounds (**F1, G1**) and replicas (**F2, G2**) in engulfment (**F**) and degradation (**G**). For variability between compounds, bars in the upper row represent the mean of normalized mean values for all compounds, and error bars represent standard deviation. For variability between replicas, bars in the lower row represent the mean standard deviation for all compounds, and error bars represent the standard deviation of standard deviations. **H, I**) Effect that the elimination of toxic compounds has on variability between compounds (**H1, I1**) and between replicas (**H2, I2**) in engulfment (**H**) and degradation (**I**). **** represents P value < 0.0001 in a Dunn´s test, after a Kruskal-Wallis test was significant.

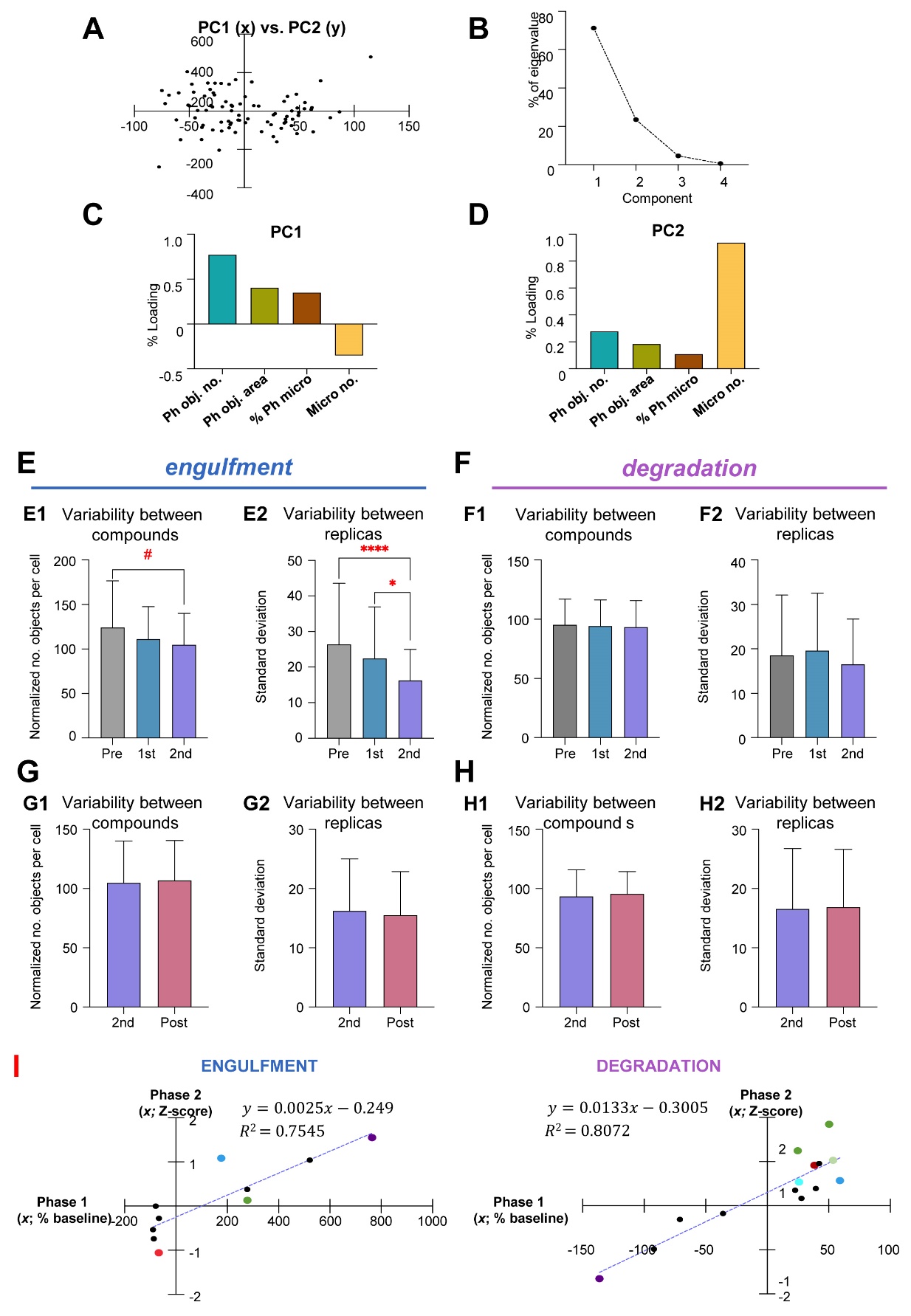
**Supplementary Figure 2.** **Compound homogenization and toxicity filtering in Phase 2**. **A**) PC1 vs PC2 dot plot representing 29 hit compounds at 3 concentrations (1, 10, and 100μM). **B**) Graphical representation of the percentage that the PC component represents on the total variance. Eigenvalues are the magnitude (the variance) associated with each PC. **C, D**) Bar graph representing the weight that each variable has on PC1 (C), or PC2 (D). **E, F**). Comparison of variability between compounds (**E1, F1**) and replicas (**E2, F2**) in engulfment € and degradation (**F**). For variability between compounds, bars in the upper row represent the mean of normalized mean values for all compounds, and error bars represent standard deviations. For variability between replicas, bars in the lower row represent the mean standard deviation for all compounds, and error bars represent the standard deviation of standard deviations. (**G, H**) Effect that the elimination of toxic compounds has on variability between compounds (**C1, D1**) and between replicas (**G2, H2**) in engulfment (**G**) and degradation (**H**). # represents P value < 0.1; * represents P value < 0.05; **** represents P value < 0.0001. **I**) Linear regression analysis comparing the effect of the 29 hit compounds in Phase 1 and Phase 2.

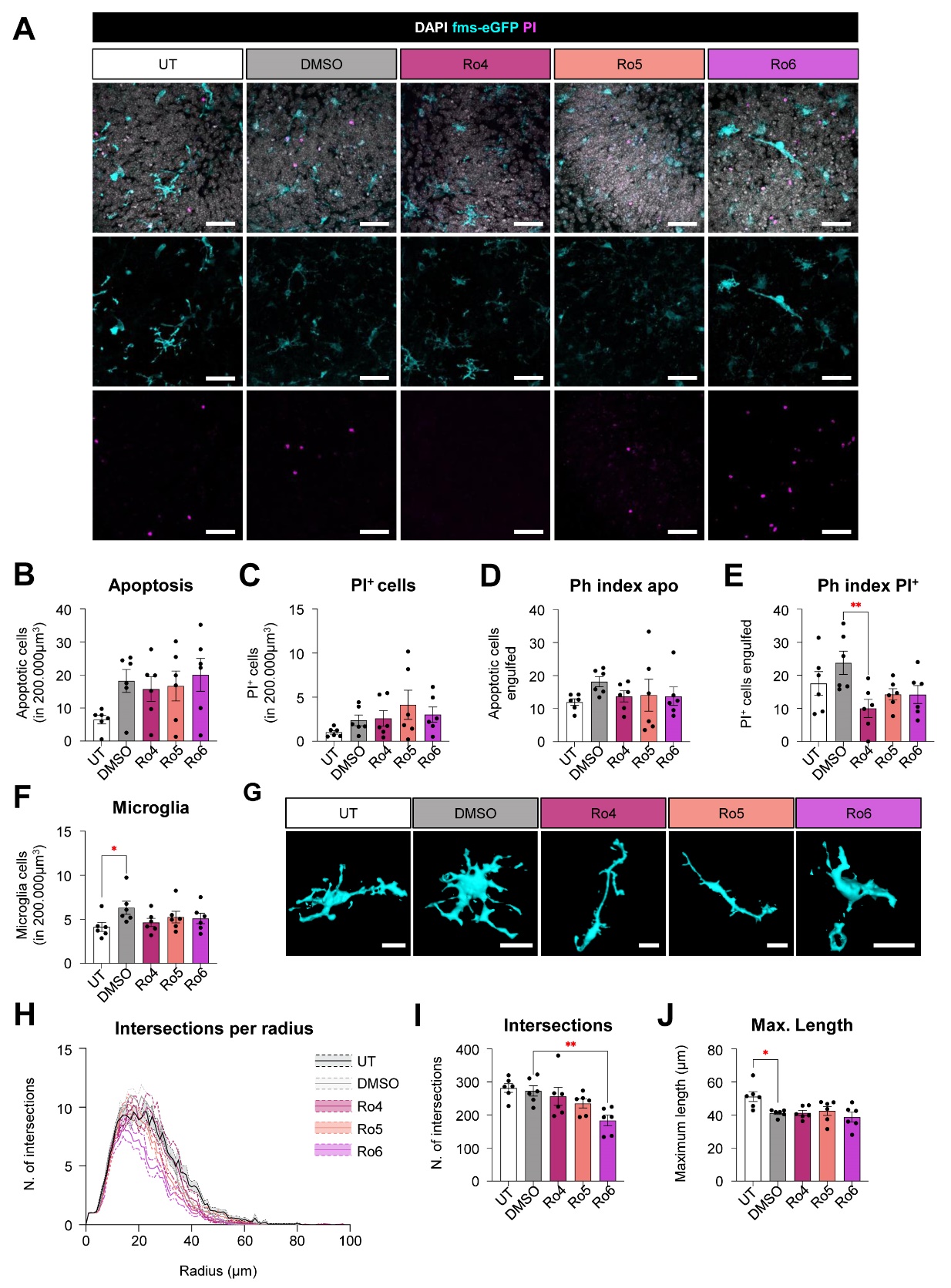
**Supplementary Figure 3. Ro4, Ro5 and Ro6 do not modulate microglial phagocytosis in organotypic slices. A**) Representative confocal z-stack of organotypic slices treated with Ro4, Ro5, and Ro6, as well as Untreated (UT) and DMSO conditions, showing cell nuclei stained with DAPI (white), microglia (cyan), and PI^+^ cells (magenta). **B, C**) Number of apoptotic and PI^+^ cells in 200.000μm^3^, respectively. **D, E**) Percentage of apoptotic and PI^+^ cells engulfed by microglia (Ph index), respectively. **F**) Number of microglia cells in 200.000μm^3^. **G**) 3D reconstructions of microglia. **H**) Histogram showing the number of microglial intersections in each radius starting from the soma. **I**) Sum of all intersections of microglial processes. **J**) Maximum length of microglial processes. Scale bars in **A** and **G**: 50μm. In **A**, z=10.5μm. Bars show mean ± SEM. n=6 (UT), n=6 (DMSO), n=6 (Ro4), n=6 (Ro5), and n=6 (Ro6), where n is the number of animals. In **D**, **E**, **I**, and **J**, data were analyzed with a one-way ANOVA followed by a Dunnett’s multiple comparisons test (*vs* DMSO). In **B**, **C**, and **F**, data were data^2^, Log10, and Log10 transformed, respectively, to comply with normality and homoscedasticity and then analyzed with a one-way ANOVA followed by a Dunnett’s multiple comparisons test. # represents p < 0.1; * represents p < 0.05; ** represents p < 0.01.

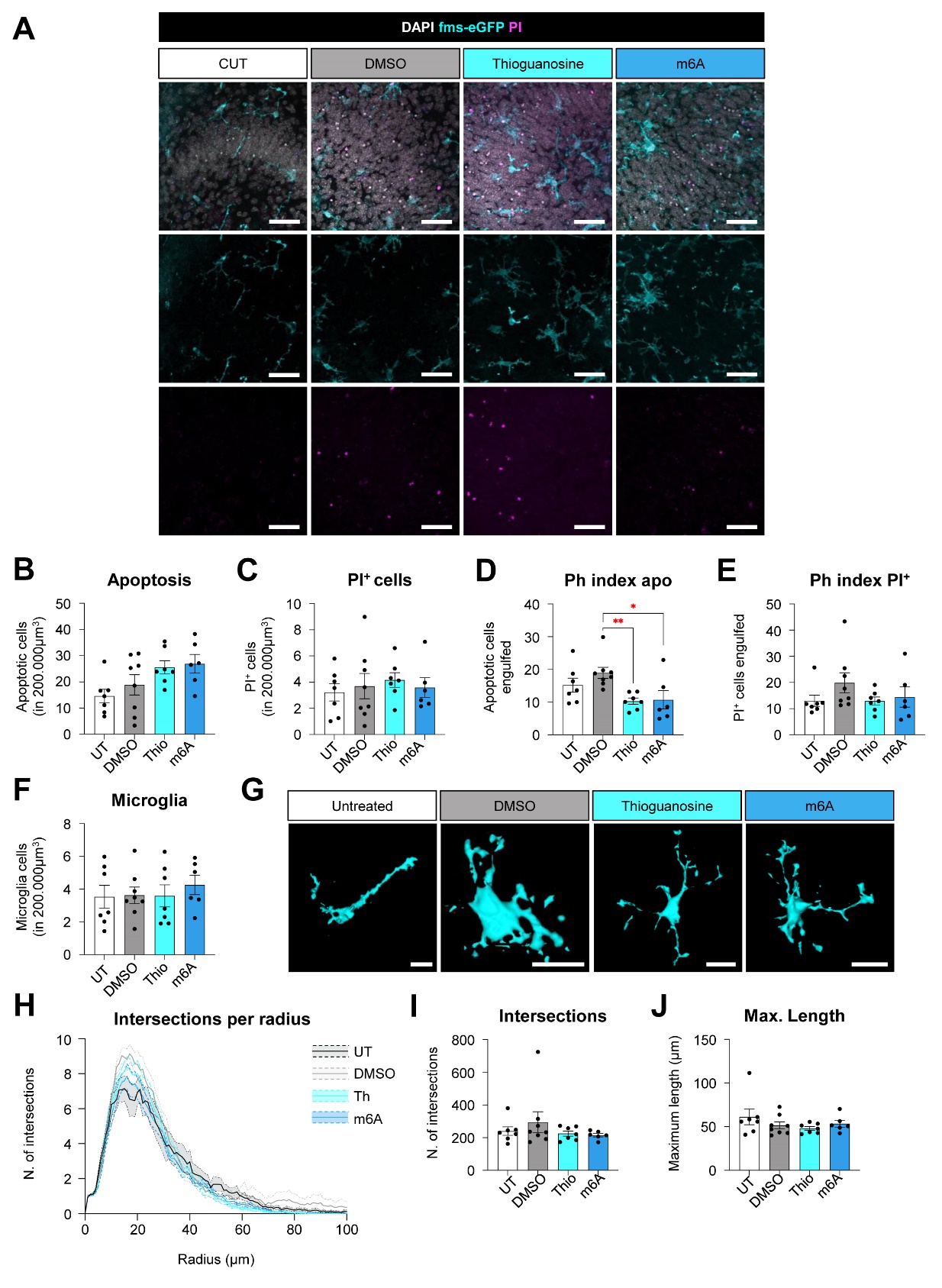
**Supplementary Figure 4**. **Thioguanosine and m6A inhibit microglial phagocytosis in organotypic slices. A**) Representative confocal z-stack of organotypic slices treated with thioguanosine (Thio) and N6-methyladenosine (m6A), as well as Untreated (UT) and DMSO conditions, showing cell nuclei stained with DAPI (white), microglia (cyan), and PI^+^ cells (magenta). **B, C**) Number of apoptotic and PI^+^ cells in 200.000μm^3^, respectively. **D, E**) Percentage of apoptotic and PI^+^ cells engulfed by microglia (Ph index), respectively. **F**) Number of microglia cells in 200.000μm^3^. **G**) 3D reconstructions of microglia. **H**) Histogram showing the number of microglial intersections in each radius starting from the soma. **I**) Sum of all intersections of microglial processes. **J**) Maximum length of microglial processes. Scale bars in **A** and **G**: 50μm. In **A**, z=10.5μm. Bars show mean ± SEM. n=7 (UT), n=8 (DMSO), n=7 (Thio), and n=6 (m6A), where n is the number of animals. In **B**, **C**, **D**, and **F**, data were analyzed with a one-way ANOVA followed by a Dunnett’s multiple comparisons test (*vs* DMSO). In **E**, **I**, and **J**, data were Log10, 1/data, and 1/data transformed, respectively, to comply with normality and homoscedasticity and then analyzed with a one-way ANOVA followed by a Dunnett’s multiple comparisons test. # represents p < 0.1; * represents p < 0.05; * represents p < 0.01.

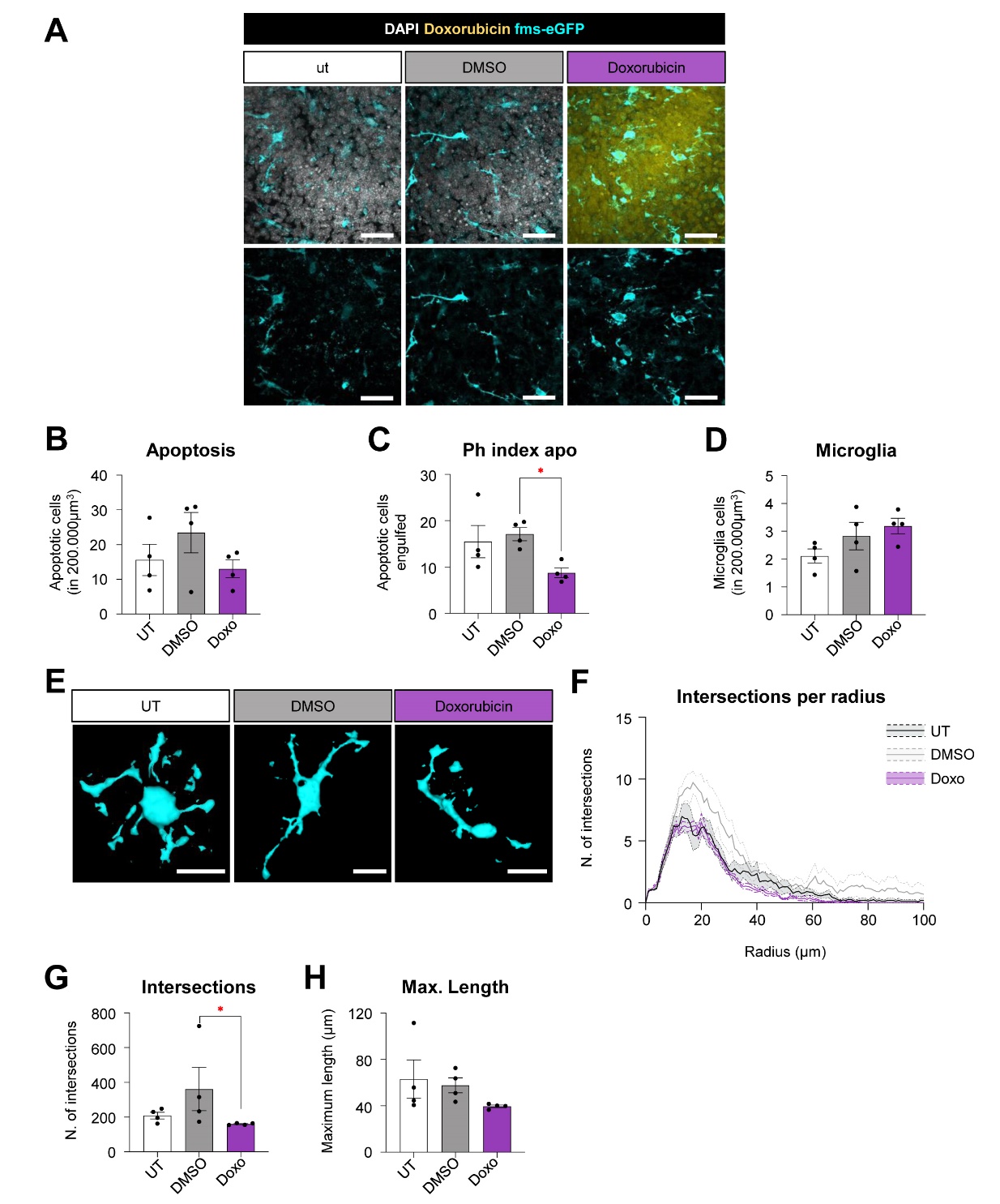
**Supplementary Figure 5. Doxorubicin inhibits microglial phagocytosis in organotypic slices. A**) Representative confocal z-stack of organotypic slices treated with doxorubicin (Doxo), as well as Untreated (UT) and DMSO conditions, showing cell nuclei stained with DAPI (white) and microglia (cyan). In doxorubicin-treated slices, cell nuclei are stained with doxorubicin (yellow). **B**) Number of apoptotic cells in 200.000μm^3^. **C**) Percentage of apoptotic cells engulfed by microglia (Ph index). **D**) Number of microglia cells in 200.000μm^3^. **E**) 3D reconstructions of microglia. **F**) Histogram showing the number of intersections in each radius starting from the soma. **G**) Sum of all intersections of microglial processes. **H**) Maximum length of microglial processes in each condition. Scale bars in **A** and **E**: 50μm. In **A**, z=10.5μm. Bars show mean ± SEM. n=4 (UT), n=4 (DMSO), and n=4 (Doxo), where n is the number of animals. In **B-D** and **H**, data were analyzed with a one-way ANOVA followed by a Dunnett’s multiple comparisons test (*vs* DMSO). In **G**, data were 1/data transformed to comply with normality and homoscedasticity and then analyzed with a one-way ANOVA followed by a Dunnett’s multiple comparisons test. # represents p < 0.1; * represents p < 0.05.

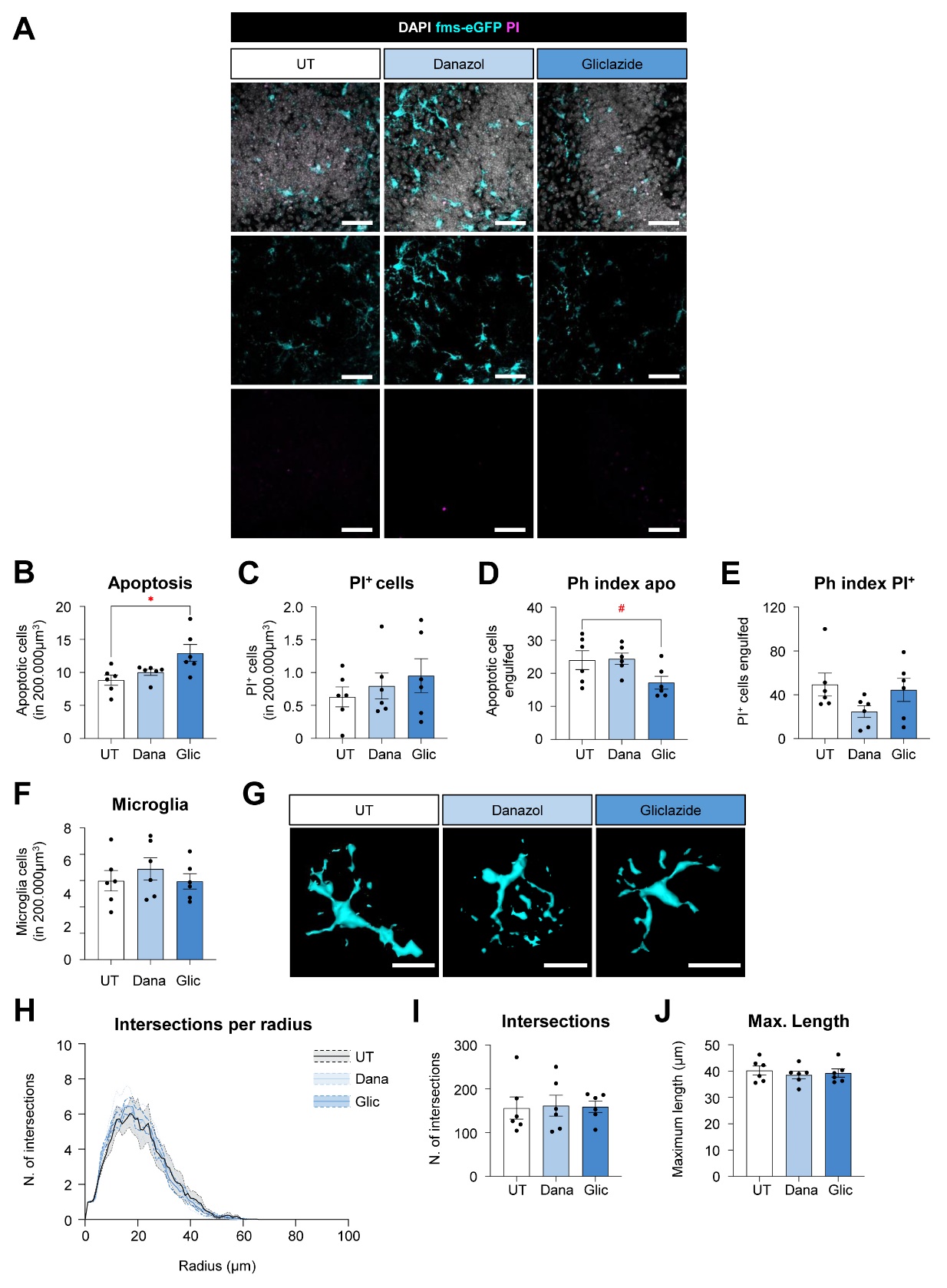
**Supplementary Figure 6**. **Gliclazide but not danazol, inhibits microglial phagocytosis in organotypic slices. A**) Representative confocal z-stack of organotypic slices treated with danazol (Dana) and Gliclazide (Glic), as well as Untreated (UT) conditions, showing cell nuclei stained with DAPI (white), microglia (cyan), and PI^+^ cells (magenta). **B, C**) Number of apoptotic and PI^+^ cells in 200.000μm^3^, respectively. **D, E**) Percentage of apoptotic and PI^+^ cells engulfed by microglia (Ph index), respectively. **F**) Number of microglia cells in 200.000μm^3^. **G**) 3D reconstructions of microglia. **H**) Histogram showing the number of microglial intersections in each radius starting from the soma. **I**) Sum of all intersections of microglial processes. **J**) Maximum length of microglial processes. Scale bars in **A** and **G**: 50μm. In **A**, z=10.5μm. Bars show mean ± SEM. n=6 (UT), n=6 (Dana), and n=6 (Glic), where n is the number of animals. In **B-J**, data were analyzed with a one-way ANOVA followed by a Dunnett’s multiple comparisons test (*vs* Untreated). # represents p < 0.1; * represents p < 0.05.

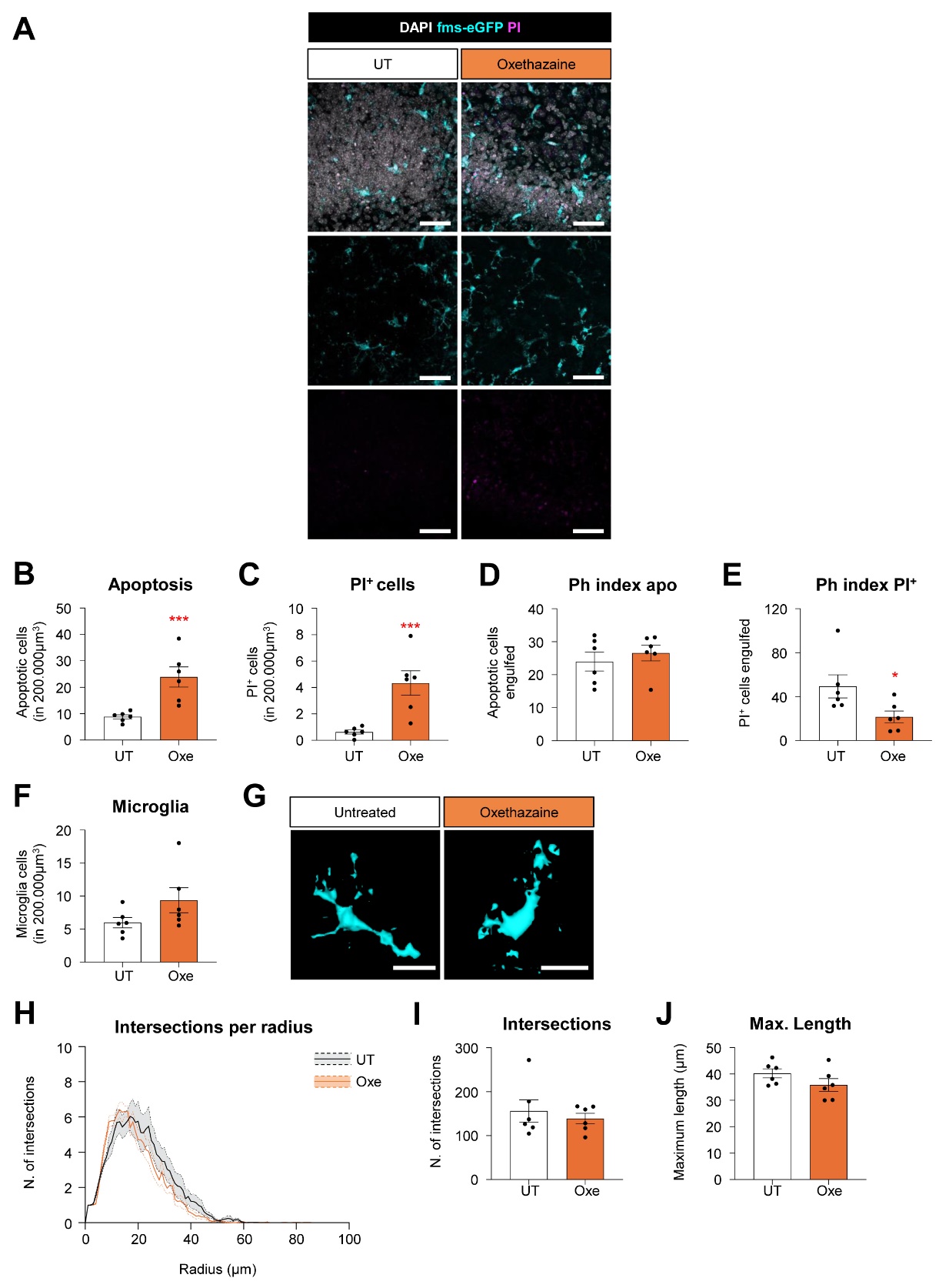
**Supplementary Figure 7. Oxethazaine is toxic in organotypic slices. A**) Representative confocal z-stack of organotypic slices treated with oxethazaine (Oxe), as well as Untreated (UT) conditions, showing cell nuclei stained with DAPI (white), microglia (cyan), and PI^+^ cells (magenta). **B, C**) Number of apoptotic and PI^+^ cells in 200.000μm^3^, respectively. **D, E**) Percentage of apoptotic and PI^+^ cells engulfed by microglia (Ph index), respectively. **F**) Number of microglia cells in 200.000μm^3^. **G**) 3D reconstructions of microglia. **H**) Histogram showing the number of microglial intersections in each radius starting from the soma. **I**) Sum of all intersections of microglial processes. **J**) Maximum length of microglial processes. Scale bars in **A** and **G**: 50μm. In **A**, z=10.5μm. Bars show mean ± SEM. n=6 (UT) and n=6 (Oxe), where n is the number of animals. In **D-J**, data were analyzed with an unpaired t-test. In **B-C**, data were Log10 and SQRT transformed to comply with normality and homoscedasticity and then analyzed with an unpaired t-test. * represents p < 0.05; *** represents p < 0.001.

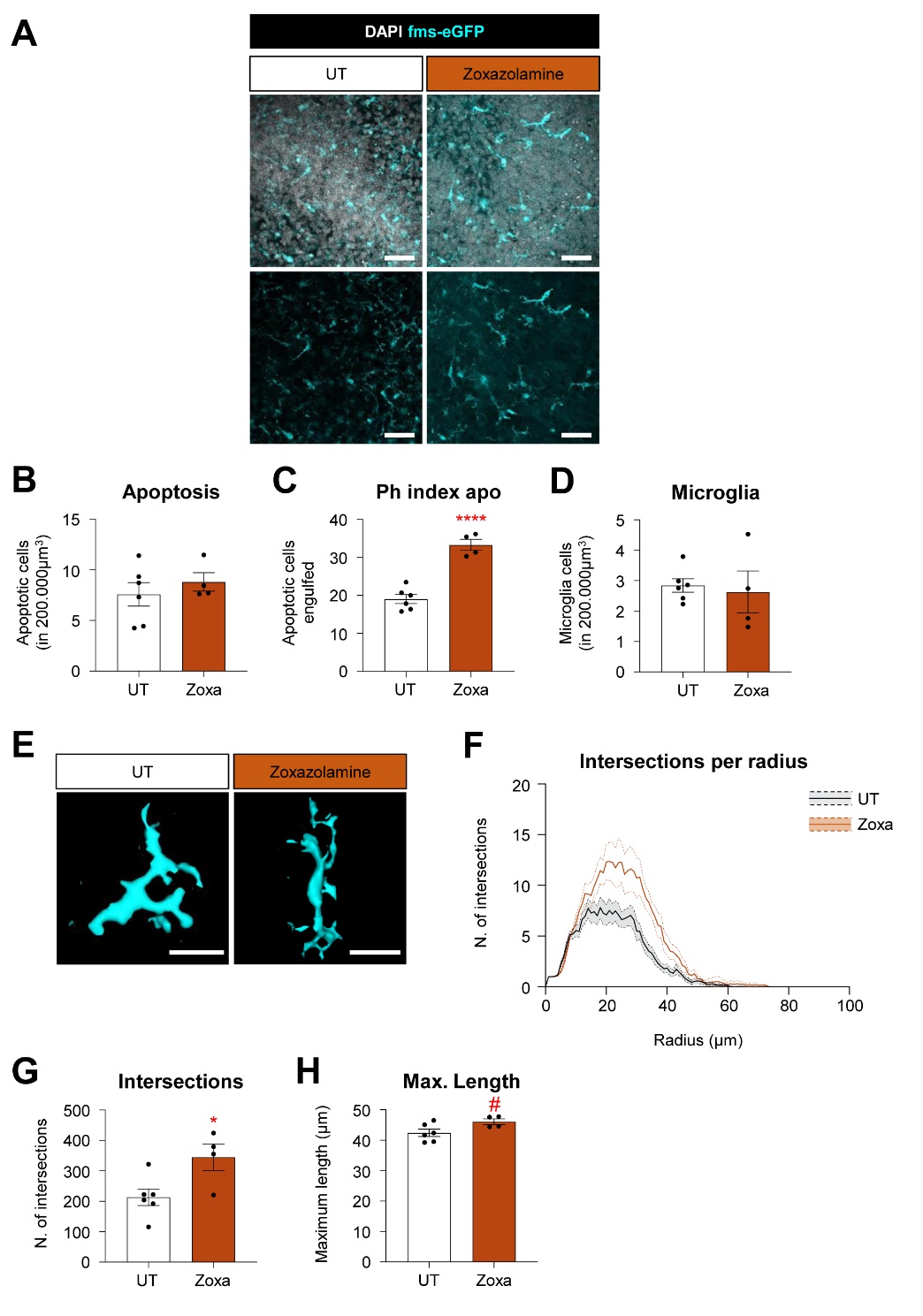

**Supplementary Figure 8. Zoxazolamine is a potential phagocytosis enhancer in organotypic slices. A**) Representative confocal z-stack of organotypic slices treated with zoxazolamine (Z), as well as Untreated (UT) conditions, showing cell nuclei stained with DAPI (white) and microglia (cyan). **B**) Number of apoptotic in 200.000μm^3^. **C**) Percentage of apoptotic cells engulfed by microglia (Ph index), respectively. **D**) Number of microglia cells in 200.000μm^3^. **E**) 3D reconstructions of microglia. **F**) Histogram showing the number of microglial intersections in each radius starting from the soma. **G**) Sum of all intersections of microglial processes. **H**) Maximum length of microglial processes. Scale bars in **A** and **E**: 50μm. In **A**, z=10.5μm. Bars show mean ± SEM. n=6 (UT) and n=4 (Zoxa), where n is the number of animals. In **B-H**, data were analyzed with an unpaired t-test. # represents p < 0.1; * represents p < 0.05; **** represents p < 0.0001.

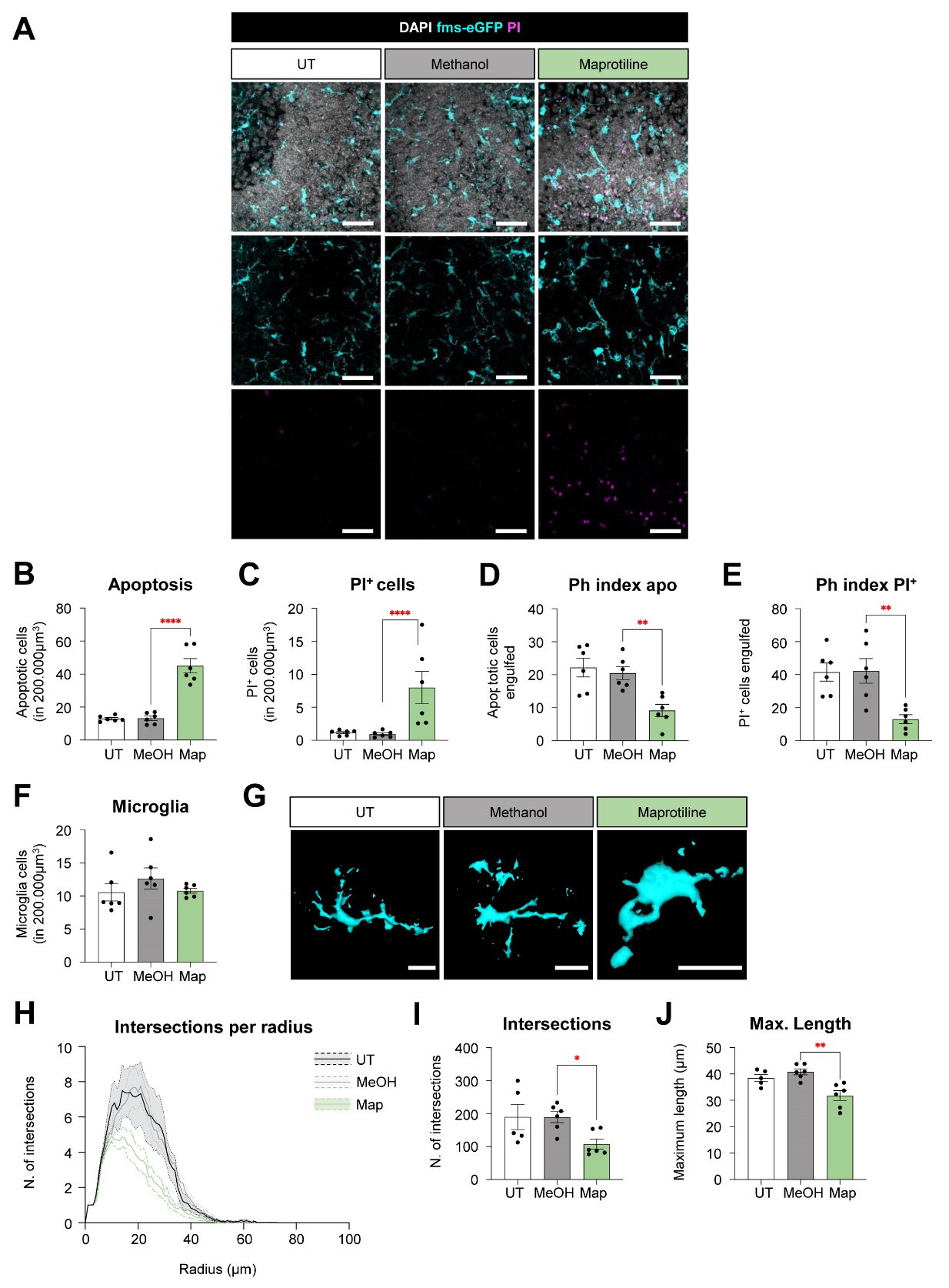
**Supplementary Figure 9. Maprotiline inhibits microglial phagocytosis in organotypic slices. A**) Representative confocal z-stack of organotypic slices treated with maprotiline (M), as well as Untreated (UT) and Methanol (MetOH) conditions, showing cell nuclei stained with DAPI (white), microglia (cyan), and PI^+^ cells (magenta). **B, C**) Number of apoptotic and PI^+^ cells in 200.000μm^3^, respectively. **D, E**) Percentage of apoptotic and PI^+^ cells engulfed by microglia (Ph index), respectively. **F**) Number of microglia cells in 200.000μm^3^. **G**) 3D reconstructions of microglia. **H**) Histogram showing the number of microglial intersections in each radius starting from the soma. **I**) Sum of all intersections of microglial processes. **J**) Maximum length of microglial processes. Scale bars in **A** and **G**: 50μm. In **A**, z=10.5μm. Bars show mean ± SEM. n=6 (UT), n=6 (MeOH), and n=6 (Map), where n is the number of animals. In **D-J**, data were analyzed with a one-way ANOVA followed by a Dunnett’s multiple comparisons test (*vs* Methanol). In **B-C**, data were Log10 transformed to comply with normality and homoscedasticity and then analyzed with a one-way ANOVA followed by a Dunnett’s multiple comparisons test (*vs* Methanol). * represents p < 0.05; ** represents p < 0.01; *** represents p < 0.001; **** represents p < 0.0001.

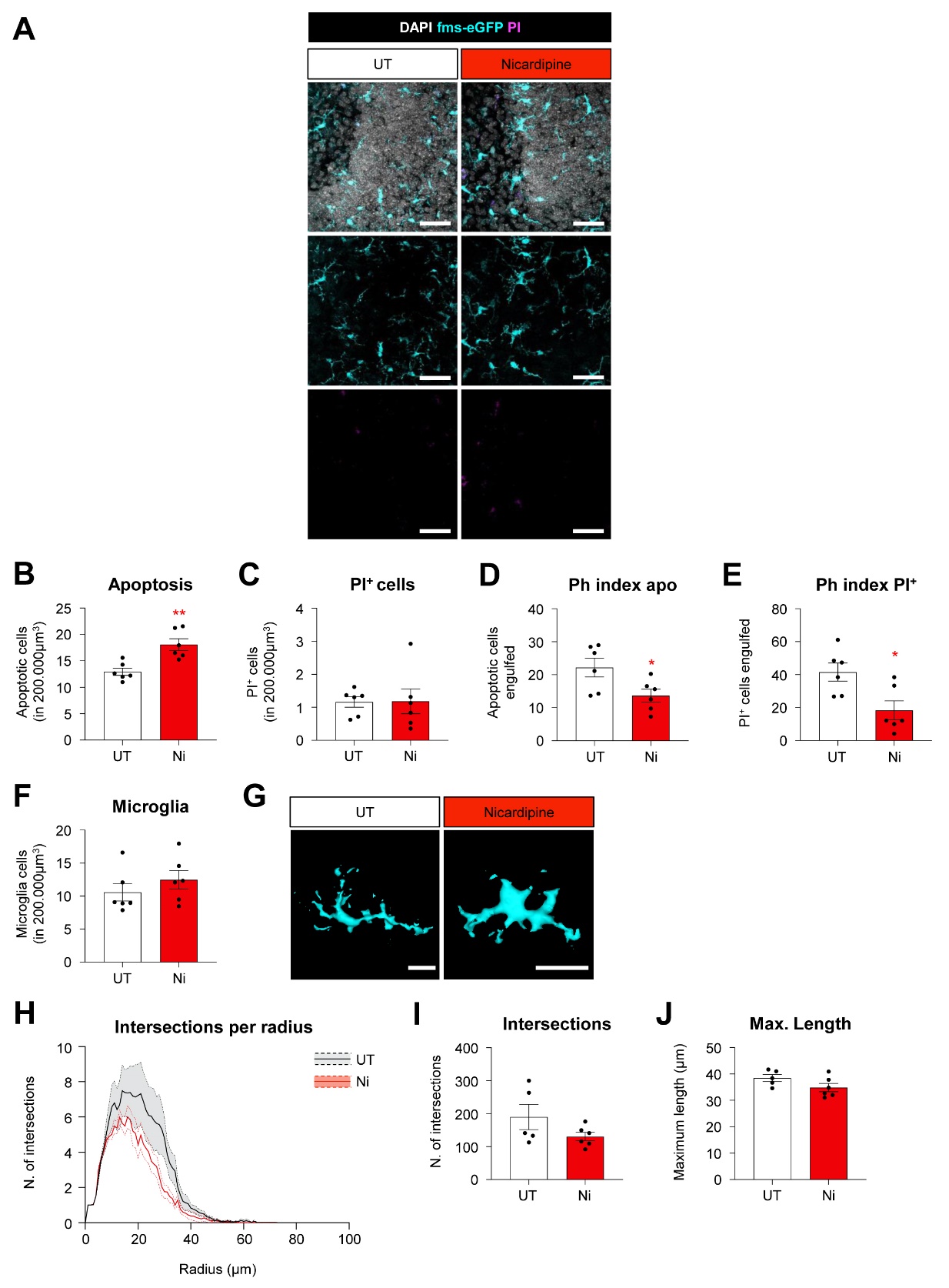
**Supplementary Figure 10. Nicardipine inhibits microglial phagocytosis in organotypic slices. A**) Representative confocal z-stack of organotypic slices treated with nicardipine (Ni), as well as Untreated (UT) conditions, showing cell nuclei stained with DAPI (white), microglia (cyan), and PI^+^ cells (magenta). **B, C**) Number of apoptotic and PI^+^ cells in 200.000μm^3^, respectively. **D, E**) Percentage of apoptotic and PI^+^ cells engulfed by microglia (Ph index), respectively. **F**) Number of microglia cells in 200.000μm^3^. **G**) 3D reconstructions of microglia. **H**) Histogram showing the number of microglial intersections in each radius starting from the soma. **I**) Sum of all intersections of microglial processes. **J**) Maximum length of microglial processes. Scale bars in **A** and **G**: 50μm. In **A**, z=10.5μm. Bars show mean ± SEM. n=5-6 (UT) and n=6 (Ni), where n is the number of animals. In **C-F** and **J**, data were analyzed with an unpaired t-test. In **I**, data were Log10 transformed to comply with normality and homoscedasticity and then analyzed with an unpaired t-test. * represents p < 0.05; ** represents p < 0.01.

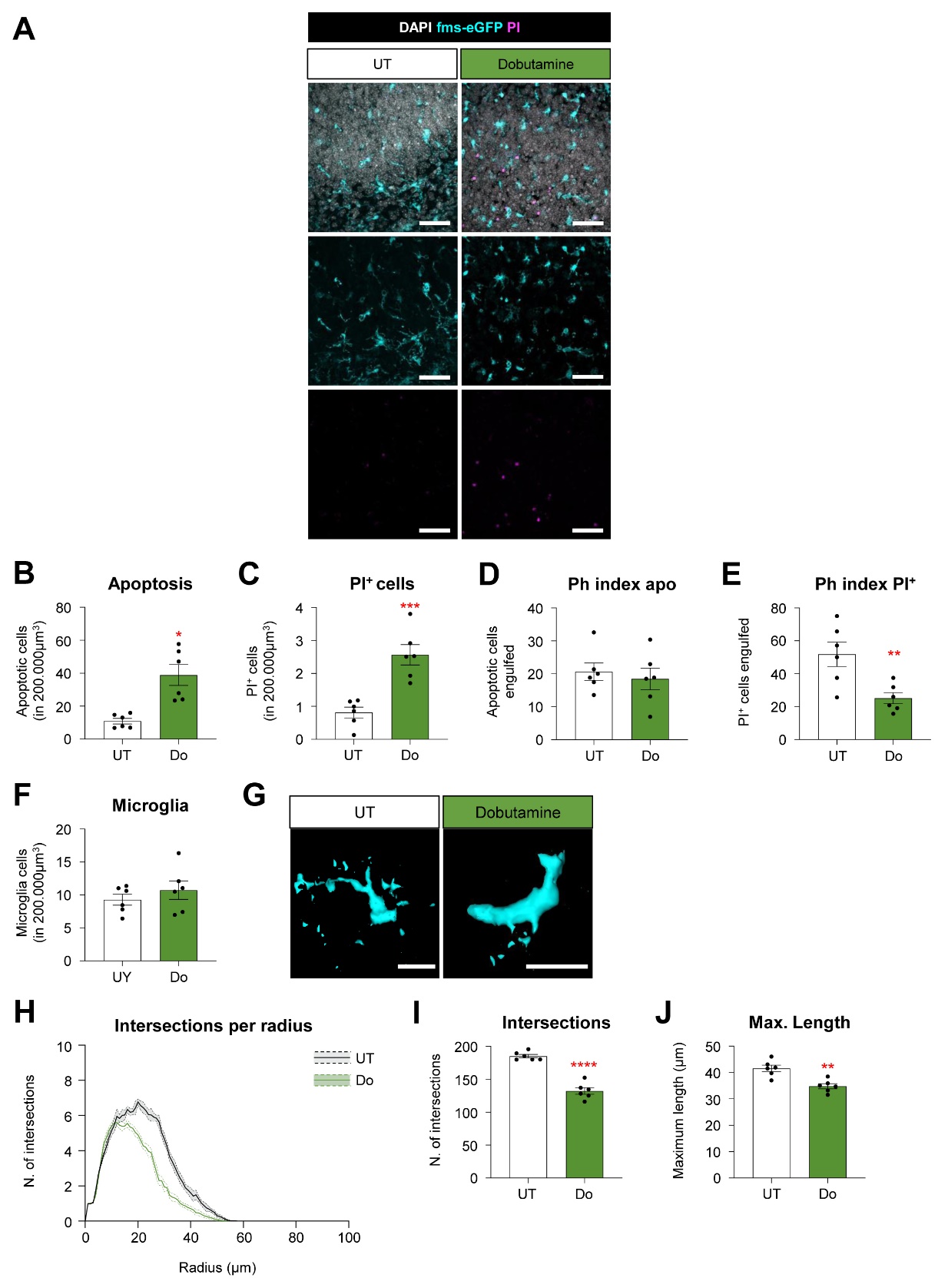
**Supplementary Figure 11. Dobutamine is toxic in organotypic slices. A**) Representative confocal z-stack of organotypic slices treated with Dobutamine (Do), as well as Untreated (UT) conditions, showing cell nuclei stained with DAPI (white), microglia (cyan), and PI^+^ cells (magenta). **B, C**) Number of apoptotic and PI^+^ cells in 200.000μm^3^, respectively. **D, E**) Percentage of apoptotic and PI^+^ cells engulfed by microglia (Ph index), respectively. **F**) Number of microglia cells in 200.000μm^3^. **G**) 3D reconstructions of microglia. **H**) Histogram showing the number of microglial intersections in each radius starting from the soma. **I**) Sum of all intersections of microglial processes. **J**) Maximum length of microglial processes. Scale bars in **A** and **G**: 50μm. In **A**, z=10.5μm. Bars show mean ± SEM. n=6 (UT) and n=6 (Do), where n is the number of animals. In **C-J**, data were analyzed with an unpaired t-test. In **B**, data were SQRT transformed to comply with normality and homoscedasticity and then analyzed with an unpaired t-test. * represents p < 0.05; ** represents p < 0.01; *** represents p < 0.001; **** represents p < 0.0001.
